# Pollen morphology of Polish species from the genus *Rubus* L. (Rosaceae) and its systematic importance

**DOI:** 10.1101/734202

**Authors:** Kacper Lechowicz, Dorota Wrońska-Pilarek, Jan Bocianowski, Tomasz Maliński

## Abstract

The genus *Rubus* L. (Rosaceae) has as yet not been investigated satisfactorily in terms of palynology. This genus is taxonomically very difficult due to the large number of species and problems with their delimitation, as well as very different distribution areas of particular species. The aim of this study was to investigate pollen morphology and for the first time the ranges of intrageneric and interspecific variability of *Rubus* species, as well as verify the taxonomic usefulness of these traits in distinguishing studied taxa from this genus. They were analysed for 11 quantitative pollen characteristics and the following qualitative ones: exine ornamentation, pollen outline and shape, as well as bridge structure. Analyses were conducted on a total of 1740 pollen grains, which represent 58 blackberry species belonging to a majority of subgenera and all the sections and series found in Poland. The diagnostic characters included exine ornamentation (exine ornamentation type, width and direction of grooves and striae, number and diameter of perforations) and length of the polar axis (P). The arrangement of the examined species on the dendrogram does not corroborate division of the genus *Rubus* into subgenera, sections and series currently adopted in taxonomy. The lack of dependence may result from apomixis observed in *Rubus*, which could reduce natural variability. Pollen features should be treated in taxonomy as auxiliary, because they fail to differentiate several (10) individual species, while the other ones create groups with similar pollen traits.

## Introduction

*Rubus* L. is a large and diverse genus in the Rosaceae family with a worldwide distribution, including hundreds or even thousands of published species names and infrageneric taxa [1, 2]. Depending on which classification you follow, historic or modern, the number of *Rubus* species may vary from 429 to 750 or up to 1000 worldwide [3–9].

The genus *Rubus* L. belongs to the tribe *Rubeae* Dumort., subfamily *Rosoideae*, family *Rosaceae* Juss., order *Rosales* Perleb and clade *Rosidae* Takht. [10, 11]. According to the new APG IV classification, the studied genus belongs to the clades Superrosids, Rosids and the order Rosales [12]. The genus *Rubus* was traditionally divided into 12 subgenera [13, 14]. The current classification recognises 13 subgenera, with the largest subgenus *Rubus* in turn divided into 12 sections [11]. However, this classification is clearly arbitrary, as many of the subgenera have been shown to be poly- or paraphyletic [15]. Most of the European blackberries belong to the typical subgenus – *Rubus*. Other subgenera were also distinguished from it: *Chamaerubus, Cylactis, Anoplobatus* and *Idaeobatus*, which were represented by individual species [9, 16].

According to Weber [9], about 250 to 300 species of brambles are found in Central and North-Western Europe. In turn, Stace [17] described approx. 300 species from the British Isles alone. In Poland, the occurrence of 108 species from the genus *Rubus* has been confirmed so far [18]. Since the publication of the genus *Rubus* monograph written by the most outstanding Polish batologist, prof. Jerzy Zieliński [16], five new bramble species have been described in Poland and 10 new species for the Polish flora have been recorded [18]. Although brambles have been a group of plants widespread throughout Europe, their phytogeographic, ecological and genetic diagnosis is still incomplete.

The genus *Rubus* is a highly complex one, particularly the subgenus *Rubus*, with poliploidy hybridisation and apparently frequent facultative apomixis, thus leading to great variation in the subgenus and making species classification one of the grand challenges of systematic botany [9, 16, 19]. Apomixis is characteristic almost exclusively to the subgenus *Rubus*, embracing most of the European bramble species. Apomixis in brambles gives rise to grains that are mature and of typical structure, as well as much smaller and not fully developed pollen. Facultative apomicts produce fewer undeveloped grains (several per cent) than obligate ones, in which they constitute from 10 to 25% [20].

Because pollen grains have an unique biological characteristics, contain a large amount of genetic information, and exhibit strong genetic conservation, they can be used for species identification [21–23]. Due to considerable difficulties in recognising particular blackberry species, pollen grains of most blackberry species have not been described in the palynological literature so far. To date only a few authors have studied pollen morphology of European taxa from this critical genus, and they are mostly older works, in which only several selected species (from 3 to 18) or the most important pollen grain features (pollen shape and exine ornamentation) were described. As a result, pollen grains of only 48 European bramble species have been described [24–34]. Among the 108 Polish brambles species, pollen of just 15 species has been characterised so far, of which six are endemic species [31, 32, 35].

The most important characteristics of bramble pollen grains include exine ornamentation (ornamentation type, width and orientation of striae and grooves), lenght of colpori, type of the bridge (clamped vs. stretched), costae colpi and the number and size of perforations [25–32, 34–48]. According to Tomlik-Wyremblewska [31, 42], pollen size and shape prove to be poor criteria in species identification.

Despite relatively numerous publications, our knowledge concerning blackberry pollen morphology is far from complete, because the available descriptions are usually brief and sometimes limited to mean dimensions. Moreover, researchers typically analyse individual, most important pollen grain characters (such as pollen size and exine ornamentation); alternatively, only some selected species were characterised. Therefore, the aim of the presented study was to perform a comprehensive analysis of relationships among the species within the taxonomically challenging genus *Rubus* L., based on pollen features of 58 species, representing four subgenera, all three sections and 23 series found in Poland. Many of the studied blackberry species are distributed throughout Europe. Another aim of this study was to discuss the taxonomic significance of pollen morphology with reference to the current classification of this genus according to Zieliński [16]. In addition, the intrageneric and interspecific variability of pollen grains in the *Rubus* species under investigation has not yet been comprehensively analysed.

## Materials and methods

### Pollen morphology

The study was conducted on 58 Polish and European *Rubus* species, which represent four out of five subgenera, all three sections and all 23 series of brambles found in Poland, including all six Polish endemic species (*R. capitulatus, R. chaerophylloides, R. ostroviensis, R. posnaniensis, R. seebergensis* and *R. spribillei*). We also included four common in Poland, alien blackberries species (*R. allegheniensis, R. canadensis, R. odoratus, R. xanthocarpus*) from Asia and North America. A list of the species analysed with their affiliation to particular taxa is shown in Table 1.

**Table 1.**
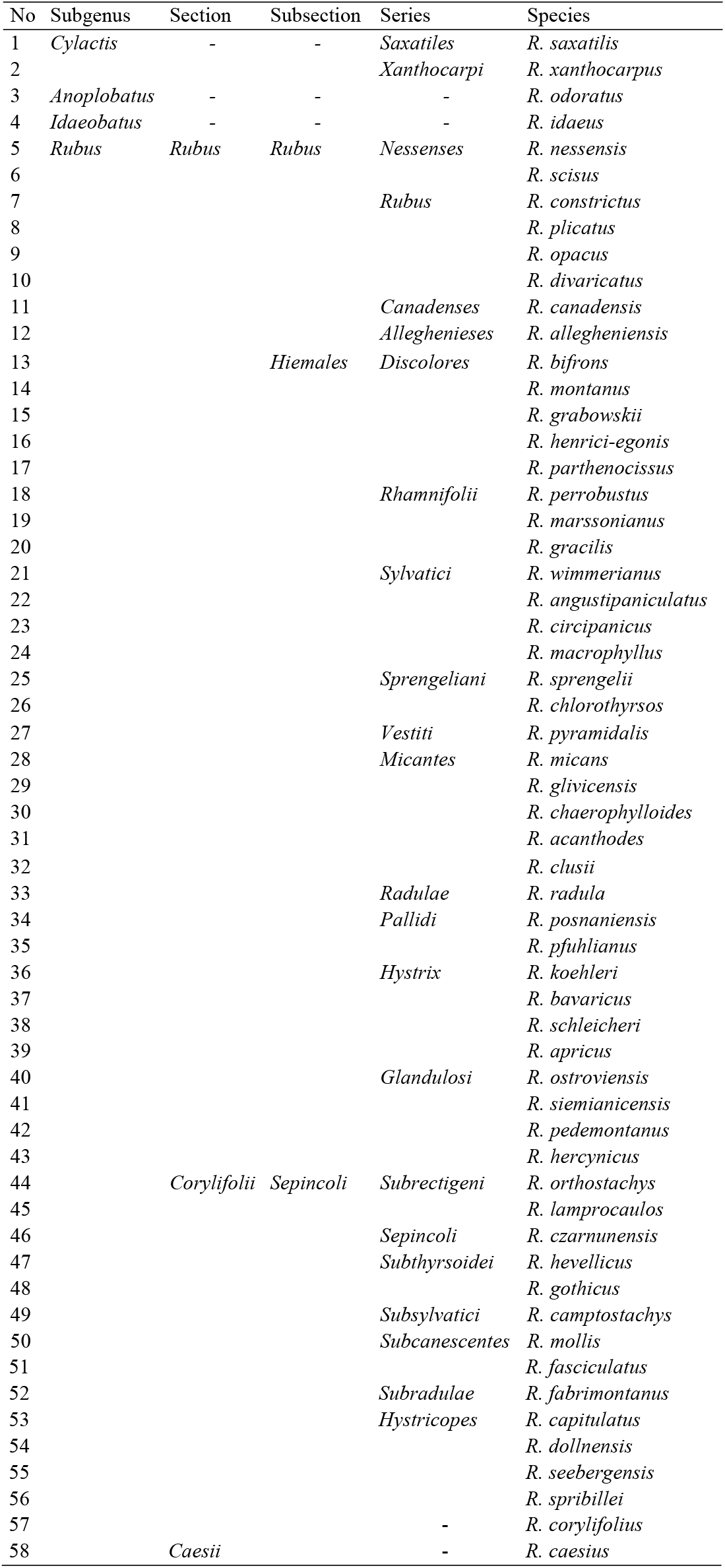
The taxonomic classification of the *Rubus* species studied.

In this paper, the taxonomic classification of the studied taxa from the genus *Rubus* was adopted from Zieliński [16], with further modifications [18]. The verification of the taxa was made by Prof. Jerzy Zieliński (Institute of Dendrology, Polish Academy of Sciences in Kórnik), an outstanding batologist – taxonomist specialising in the genus *Rubus*.

Several, randomly selected inflorescences (flowers) were collected from 58 natural bramble localities in Poland (Table 2). The plant material was stored in the herbarium of the Department of Forest Botany, the Poznan University of Life Sciences (PZNF).

**Table 2.** List of localities of the *Rubus* species studie.

| No | Species | Localities | Geographical coordinates | Collector, herbarium |
| --- | --- | --- | --- | --- |
| 1 | *R. acanthodes* | Poland, Dolnośląskie, Nowe Łąki near Pielgrzymka | 51°07'06,1"N, 15°46'37,5"E | Boratyńska, Dolatowska, Tomlik, Zieliński; KOR |
| 2 | *R. allegheniensis* | Poland, Zachodniopomorskie, Łukęcin near Świnoujście | 54°02'34,9"N, 14°52'23,8"E | Boratyńska, Dolatowska, Zieliński; KOR |
| 3 | *R. angustipaniculatus* | Poland, Mazowieckie, Zakrzew near Radom | 50°26'27,3"N, 21°00'02,4"E | Maliński, Zieliński; POZNF |
| 4 | *R. apricus* | Poland, Wielkopolskie, Bachorzew near Jarocin | 51°59'39,9"N, 17°33'49,9"E | Maliński, Zieliński; POZNF |
| 5 | *R. bavaricus* | Poland, Wielkopolskie, Robczysko near Leszno | 51°48'41,4"N, 16°45'38,6"E | Danielewicz, Maliński; POZNF |
| 6 | *R. bifrons* | Poland, Podkarpackie, Łukowe near Sanok | 49°25'20,1"N, 22°14'14,1"E | Oklejewicz; KOR |
| 7 | *R. caesius* | Poland, Lubuskie, Osiecznica near Krosno Odrzańskie | 52°04'45,0"N, 15°03'11,0"E | Maliński, Zieliński; POZNF |
| 8 | *R. camptostachys* | Poland, Wielkopolskie, Raków near Kępno | 51°11'16,8"N, 18°05'54,1"E | Zieliński; KOR |
| 9 | *R. canadensis* | Poland, Dolnośląskie, Bialskie Mts. near Stronie Śląskie | 50°14'59,9"N, 16°57'45,7"E | Kosiński; KOR |
| 10 | *R. capitulatus* | Poland, Wielkopolskie, Psienie-Ostrów near Pleszew | 51°57'48,2"N, 17°45'51,5"E | Danielewicz, Maliński; POZNF |
| 11 | *R. chaerophylloides* | Poland, Wielkopolskie, Laskowo near Chodzież | 53°01'19,2"N, 17°05'45,4"E | Maliński, Zieliński; POZNF |
| 12 | *R. chlorothyrsos* | Poland, Pomorskie, Bargędzino near Łeba | 54°43'53,4"N, 17°43'19,3"E | Boratyńska, Dolatowska, Zieliński; KOR |
| 13 | *R. circipanicus* | Poland, Zachodniopomorskie, Jarosławiec near Ustka | 54°32'21,3"N, 16°32'31,6"E | Zieliński; KOR |
| 14 | *R. clusii* | Poland, Małopolskie, Dobronków near Tarnów | 49°59'28,2"N, 21°20'37,5"E | Maliński, Zieliński; POZNF |
| 15 | *R. constrictus* | Poland, Małopolskie, Lipinki near Gorlice | 49°40'20,4"N, 21°17'31,6"E | Oklejewicz; KOR |
| 16 | *R. corylifolius* | Poland, Lubuskie, Różanówka near Bytom Odrzański | 51°46'05,4"N, 15°52'29,5"E | Maliński, Zieliński; POZNF |
| 17 | *R. czarnunensis* | Poland, Pomorskie, Drzewicz, Bory Tucholskie National Park | 53°51'07,3"N, 17°34'08,4"E | Tomlik, KOR |
| 18 | *R. divaricatus* | Poland, Lubuskie, Bielawy near Bytom Odrzański | 51°46'21,3"N, 15°55'09,6"E | Maliński, Zieliński; POZNF |
| 19 | *R. dollnensis* | Poland, Dolnośląskie, Młynowiec near Stronie Śląskie | 50°16'36,1"N, 16°54'04,8"E | Kosiński, Tomaszewski, Zieliński; KOR |
| 20 | *R. fabrimontanus* | Poland, Lubuskie, Tarnów Jezierny Nowa Sól | 51°51'45,1"N, 15°59'07,7"E | Maliński, Zieliński; POZNF |
| 21 | *R. fasciculatus* | Poland, Podkarpackie, Gruszowa near Przemyśl | 49°40'57,4"N, 22°40'47,2"E | Maliński, Zieliński; POZNF |
| 22 | *R. glivicensis* | Poland, Małopolskie, Mąga near Tarnów | 50°00'09,8"N, 21°20'24,7"E | Maliński, Zieliński; POZNF |
| 23 | *R. gothicus* | Poland, Wielkopolskie, Pakówka near Bojanowo | 51°40'20,7"N, 16°46'07,9"E | Maliński, Zieliński; POZNF |
| 24 | *R. grabowskii* | Poland, Lubuskie, Tarnów Jezierny Nowa Sól | 51°51'45,1"N, 15°59'07,7"E | Maliński, Zieliński; POZNF |
| 25 | *R. gracilis* | Poland, Podkarpackie, Pod Lasem, near Rzeszów | 49°53'42,5"N, 21°35'52,1"E | Maliński, Zieliński; POZNF |
| 26 | *R. henrici-egonis* | Poland, Opolskie, Barnice near Głubczyce | 50°03'02,5"N, 17°47'38,5"E | Kosiński, Tomaszewski, Zieliński; KOR |
| 27 | *R. hercynicus* | Poland, Dolnośląskie, Stare Bogaczowice near Wałbrzych | 50°50'53,7"N, 16°11'37,4"E | Boratyński, Zieliński; KOR |
| 28 | *R. hevellicus* | Poland, Wielkopolskie, Tarce near Jarocin | 52°00'02,4"N, 17°35'26,1"E | Maliński, Zieliński; POZNF |
| 29 | *R. idaeus* | Poland, Kujawsko-Pomorskie, Brodnica near Bydgoszcz | 53°15'29,2"N, 19°23'57,9"E | Tomlik; KOR |
| 30 | *R. koehleri* | Poland, Dolnośląskie, Mirsk near Świeradów-Zdrój | 50°58'19,9"N, 15°23'08,9"E | Boratyński; KOR |
| 31 | *R. lamprocaulos* | Poland, Dolnośląskie, Serby near Głogów | 51°41'04,1"N, 16°06'42,9"E | Maliński, Zieliński; POZNF |
| 32 | *R. macrophyllus* | Poland, Dolnośląskie, Przywście near Rawicz | 51°34'37,1"N, 16°52'36,1"E | Maliński, Zieliński; POZNF |
| 33 | *R. marssonianus* | Poland, Pomorskie, near Kartuzy | 54°20'03,2"N, 18°11'50,5"E | Boratyński; KOR |
| 34 | *R. micans* | Poland, Opolskie, Wieszczyzna near Prudnik | 50°19'18,2"N, 17°34'48,4"E | Kosiński, Tomaszewski, Zieliński; KOR |
| 35 | *R. mollis* | Poland, Dolnośląskie, Łądek-Zdrój, Trzykrszyska Mt. | 50°20'54,6"N, 16°52'39,9"E | Kosiński, Tomaszewski, Zieliński; KOR |
| 36 | *R. montanus* | Poland, Dolnośląskie, Kowary near Kostrzyca | 50°47'37,5"N, 15°50'01,8"E | Zieliński; KOR |
| 37 | *R. nessensis* | Poland, Dolnośląskie, Karczmisko near Kłodzko | 50°17'56,7"N, 16°49'32,8"E | Kosiński; KOR |
| 38 | *R. odoratus* | Poland, Lubelskie, Niedzwica Duża near Lublin | 51°06'51,3"N, 22°23'16,2"E | illegible name; KOR |
| 39 | *R. opacus* | Poland, Wielkopolskie, Starkowo near Leszno | 51°58'37,7"N, 16°18'35,7"E | Zieliński; KOR |
| 40 | *R. orthostachys* | Poland, Wielkopolskie, Ostatni Grosz near Krotoszyn | 50°39'54,4"N, 17°21'18,9"E | Maliński, Zieliński; POZNF |
| 41 | *R. ostroviensis* | Poland, Wielkopolskie, Wielkopolski National Park near Poznań | 52°16'26,5"N, 16°46'50,1"E | Zieliński, Maliński; POZNF |
| 42 | *R. parthenocissus* | Poland, Podkarpackie, Koniusza near Przemyśl | 49°40'57,4"N, 22°40'47,2"E | Maliński, Zieliński; POZNF |
| 43 | *R. pedemontanus* | Poland, Dolnośląskie, Nowy Kościół near Złotoryja | 51°04'20,1"N, 15°52'05,3"E | Boratyński, Zieliński; KOR |
| 44 | *R. perrobustus* | Poland, Podkarpackie, Dudyńce near Sanok | 49°39'04,9"N, 22°04'31,9"E | Oklejewicz; KOR |
| 45 | *R. pfuhlianus* | Poland, Wielkopolskie, Mieczewo near Kórnik | 52°14'20,8"N, 17°00'27,8"E | Zieliński; KOR |
| 46 | *R. plicatus* | Poland, Lubuskie, Różanówka near Bytom Odrzański | 51°46'05,4"N, 15°52'29,5"E | Maliński, Zieliński; POZNF |
| 47 | *R. posnaniensis* | Poland, Opolskie, Szybowice near Prudnik | 50°21'09,5"N, 17°29'11,9"E | Kosiński, Tomaszewski, Zieliński; KOR |
| 48 | *R. pyramidalis* | Poland, Wielkopolskie, Chruszczyny near Ostrów Wielkopolski | 51°38'41,4"N, 17°35'42,6"E | Maliński, Zieliński; POZNF |
| 49 | *R. radula* | Poland, Podkarpackie, Hermanowa near Rzeszów | 49°56'07,4"N, 22°00'40,4"E | Maliński, Zieliński; POZNF |
| 50 | *R. saxatilis* | Sweden, Abisko Östra | 68°20'56,3"N, 18°49'43,7"E | illegible name; KOR |
| 51 | *R. schleicheri* | Poland, Wielkopolskie, Kościan | 52°05'10,7"N, 16°38'41,9"E | Maliński, Zieliński; POZNF |
| 52 | *R. scisus* | Poland, Śląskie, Rudniki near Częstochowa | 50°52'33,6"N, 19°14'28,5"E | Zieliński; KOR |
| 53 | *R. seebergensis* | Poland, Wielkopolskie, Wielkopolski National Park near Poznań | 52°16'26,5"N, 16°46'50,1"E | Danielewicz; POZNF |
| 54 | *R. siemianicensis* | Poland, Wielkopolskie, Psienie-Ostrów near Pleszew | 51°57'48,2"N, 17°45'51,5"E | Danielewicz, Maliński; POZNF |
| 55 | *R. sprengelii* | Poland, Wielkopolskie, Borownica near Zduny | 51°38'20,8"N, 17°24'23,3"E | Maliński, Zieliński; POZNF |
| 56 | *R. spribillei* | Poland, Wielkopolskie, Gądky near Kórnik | 52°18'45,4"N, 17°02'47,8"E | Zieliński; POZNF |
| 57 | *R. wimmerianus* | Poland, Podkarpackie, Gniewczyzna Łańcucka near Przeworsk | 50°06'19,5"N, 22°29'43,7"E | Oklejewicz, Zatorski; POZNF |
| 58 | *R. xanthocarpus* | Poland, Świętokrzyskie, Miedzianka near Kielce | 50°50'22,5"N, 20°22'03,3"E | Maciejczak, Bróz, Zieliński; KOR |

KOR - Herbarium of the Institute of Dendrology, Polish Academy of Sciences, Kórnik, Poland, PZNF - Herbarium of the Department of Forest Botany, Poznań University of Life Sciences.

Measurements from 30 mature, randomly selected, properly developed pollen grains were made using light microscopy (LM), with 1740 pollen grains measured in total.

Pollen grains were acetolysed according to the method of Erdtman [49]. Grains were mixed with the acetolysis solution, which consisted of nine parts acetic anhydrite and one part concentrated sulphuric acid. The mixture was then heated to boiling and kept in the water bath for 2-3 min. Samples were centrifuged in the acetolysis mixture, washed with acetic acid and centrifuged again. The pollen grain samples were then mixed with 96% alcohol and centrifuged 4 times, with processed grains subsequently divided into two groups. One half of the processed sample was immersed in an alcohol-based solution of glycerin for LM, while the other was placed in 96% ethyl alcohol in preparation for scanning electron microscopy (SEM). The SEM observations were made using a Zeiss Evo 40 and the LM measurements of acetolysed pollen grain were taken using a Biolar 2308 microscope at a magnification of 640x. Pollen grains were immersed in glycerin jelly and measured using an ocular eyepiece with a scale. Measurement results were converted into micrometers by multiplying each measurement by two.

The pollen grains were analysed for 11 quantitative characters: length of the polar axis (P) and equatorial diameter (E), length of the ectoaperture (Le), thickness of the exine along the polar axis and equatorial diameter (Exp, Exe), distance between apices of two ectocolpi (d) and P/E, Le/P, Exp/P, Exe/E, d/E (apocolpium index P.A.I) ratios. The pollen shape classes (P/E ratio) were adopted according to the classification proposed by Erdtman [50]: oblate-spheroidal (0.89-0.99), spheroidal (1.00), prolate-spheroidal (1.01-1.14), subprolate (1.15-1.33), prolate (1.34-2.00) and perprolate (>2.01). In addition, the following qualitative characters were also determined: outline, shape, operculum structure and exine ornamentation.

Exine ornamentation types (I-VI) were identified based on the classification proposed by Ueda [41]. The types and subtypes of the striate exine ornamentation were characterised by the height and width of grooves, width of striae and the number and diameter of perforations.

Descriptive palynological terminology followed Punt et al. [51] and Halbritter et al. [52].

### Statistical analysis

The normality of the distributions for the studied traits (P, E, Le, d, Exp, Exe, P/E, Le/P, d/E, Exp/P and Exe/E) was tested using Shapiro-Wilk’s normality test [53]. Multivariate analysis of variance (MANOVA) was performed on the basis of the following model using the MANOVA procedure in GenStat (18th edition): **Y**=**XT**+**E**, where: **Y** is the (*n*×*p*)-dimensional matrix of observations, *n* is the number of all observations, *p* is the number of traits (in this study *p*=11), **X** is the (*n*×*k*)-dimensional matrix of design, *k* is the number of species (in this study *k*=58), **T** is the (*k*×*p*)-dimensional matrix of unknown effects and **E** – is the (*n*×*p*)-dimensional matrix of residuals. Next, one-way analyses of variance (ANOVA) were performed in order to verify the zero hypothesis on a lack of species effect in terms of values of observed traits, for each trait independently, on the basis of the following model: *y_ij_*=*μ*+*τ_i_*+*ε_ij_*, where: *ε_ij_* is the *j*th observation of the *i*th species, *μ* is the grand mean, *τ_i_* is the effect of the *i*th species and *ε_ij_* is an error observation. The arithmetical means and standard deviations of traits were calculated. Moreover, Fisher’s least significant differences (LSDs) were also estimated at the significance level α=0.001. The relationships between observed traits were assessed on the basis of Pearson’s correlation. Results were also analysed using multivariate methods. The canonical variate analysis was applied in order to present multitrait assessment of similarity for the tested species in a lower number of dimensions with the least possible loss of information [54]. This makes it possible to illustrate variation in species in terms of all the observed traits in the graphic form. The Mahalanobis distance was suggested as a measure of “polytrait” species similarity [55], which significance was verified by means of critical value D_α_ called “the least significant distance” [56]. Mahalanobis distances were calculated for species. The differences between the analysed species were verified by cluster analysis using the nearest neighbour method and Euclidean distances [57]. All the analyses were conducted using the GenStat (18th edition) statistical software package.

## Results

### General morphological description of pollen

A description of pollen grain morphology of the *Rubus* species studied is given below and illustrated with several SEM photographs (Figs 1-3). The morphological observations for the other quantitative characters of pollen grains are summarised in Table 3.

**Fig 1.**
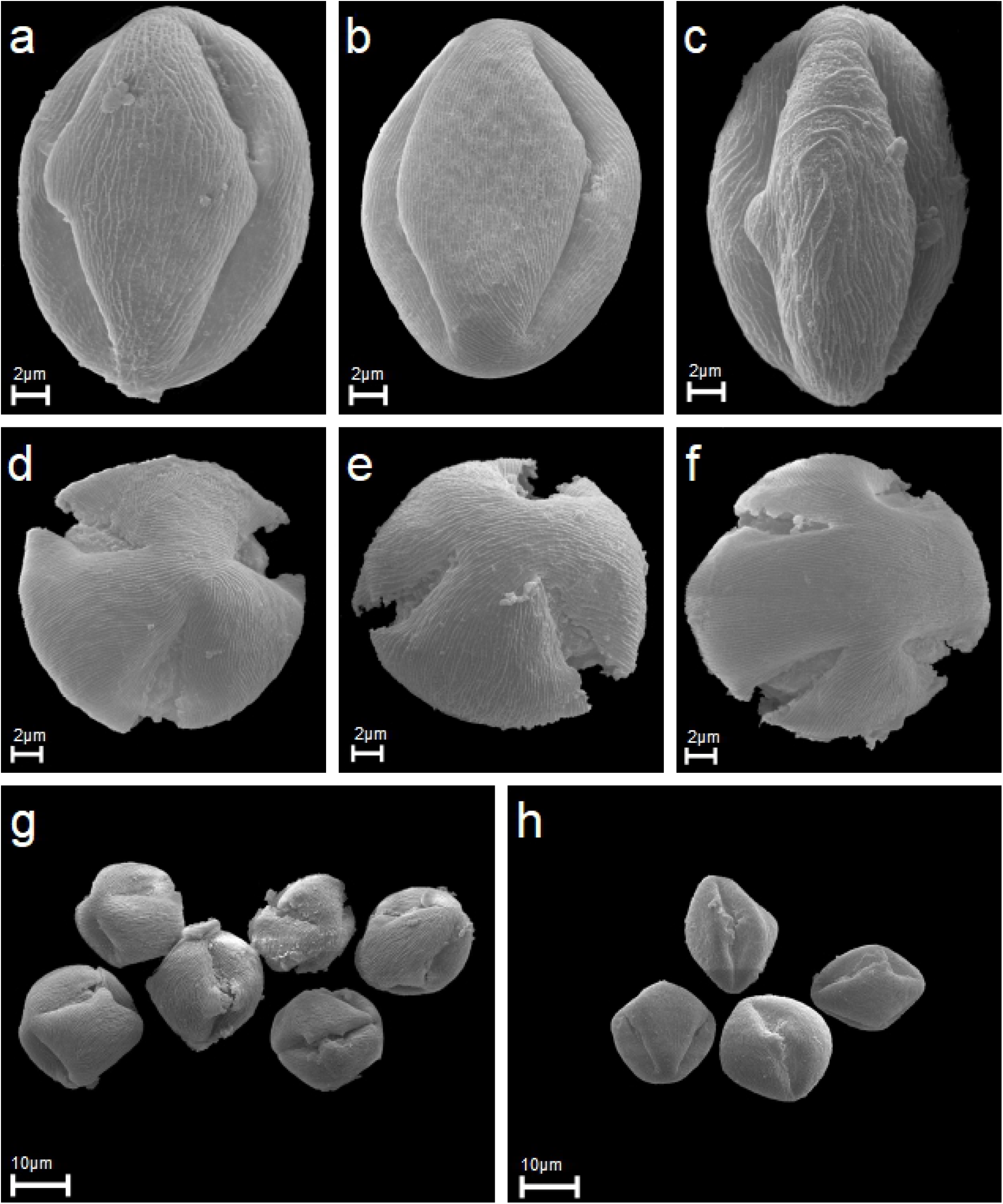
Equatorial and polar views, apertures and exine ornamentation in scanning electron microscope (SEM). (A-C) R. chlorothyrsos, R. pedemontanus, R. mollispollen grains in equatorial views, two colpori and exine ornamentation. (D-F) R. fabrimontanus, R. pfuhlianus, R. lamprocaulos pollen in polar views, three colpori and exine ornamentation. (G-H) R. angustipaniculatus, R. hevellicus six and four pollen grains in equatorial and polar views.

**Fig 2.**
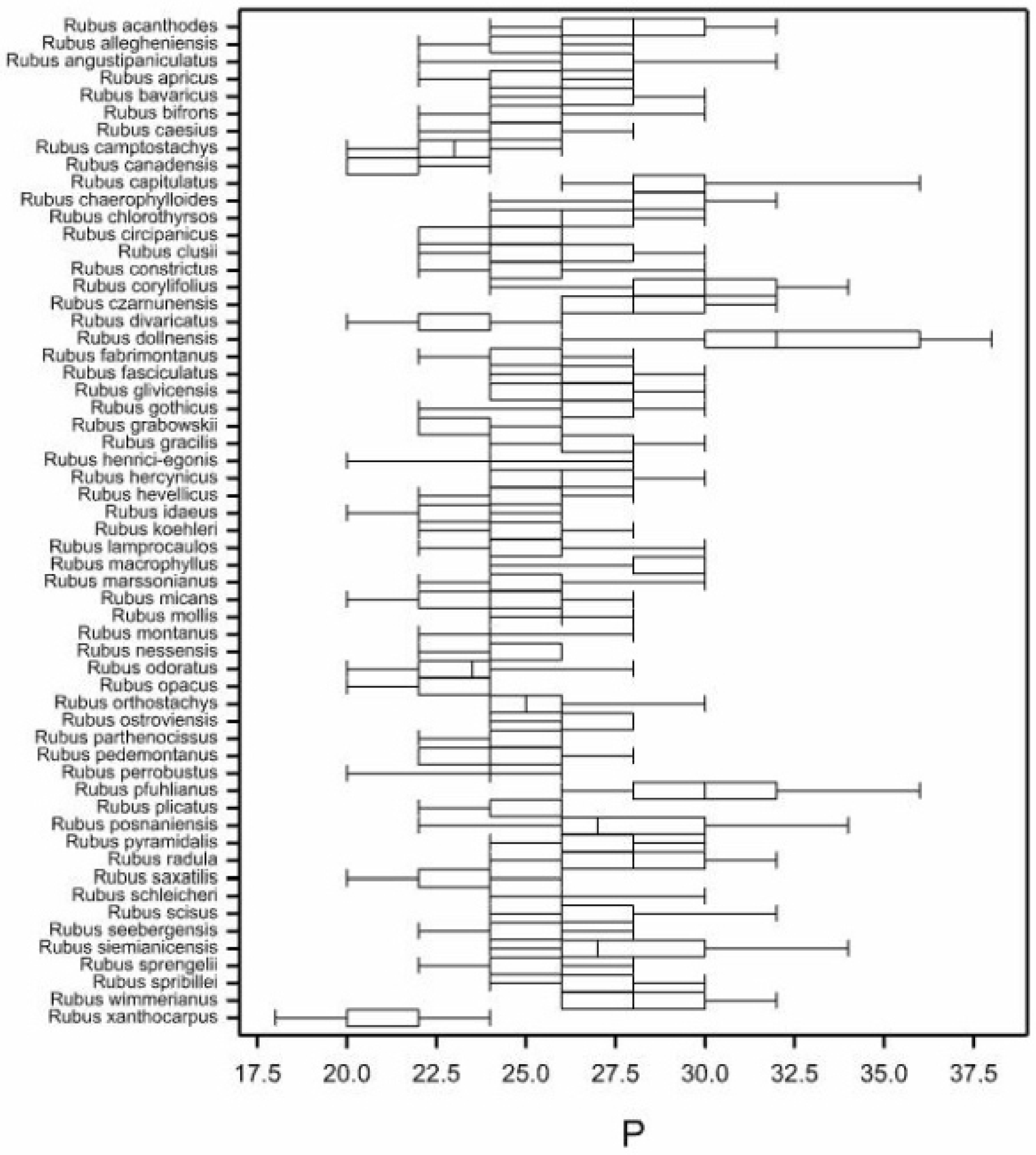
Box-and-whisker diagram of P values for 58 studied Rubus species.

**Fig 3.**
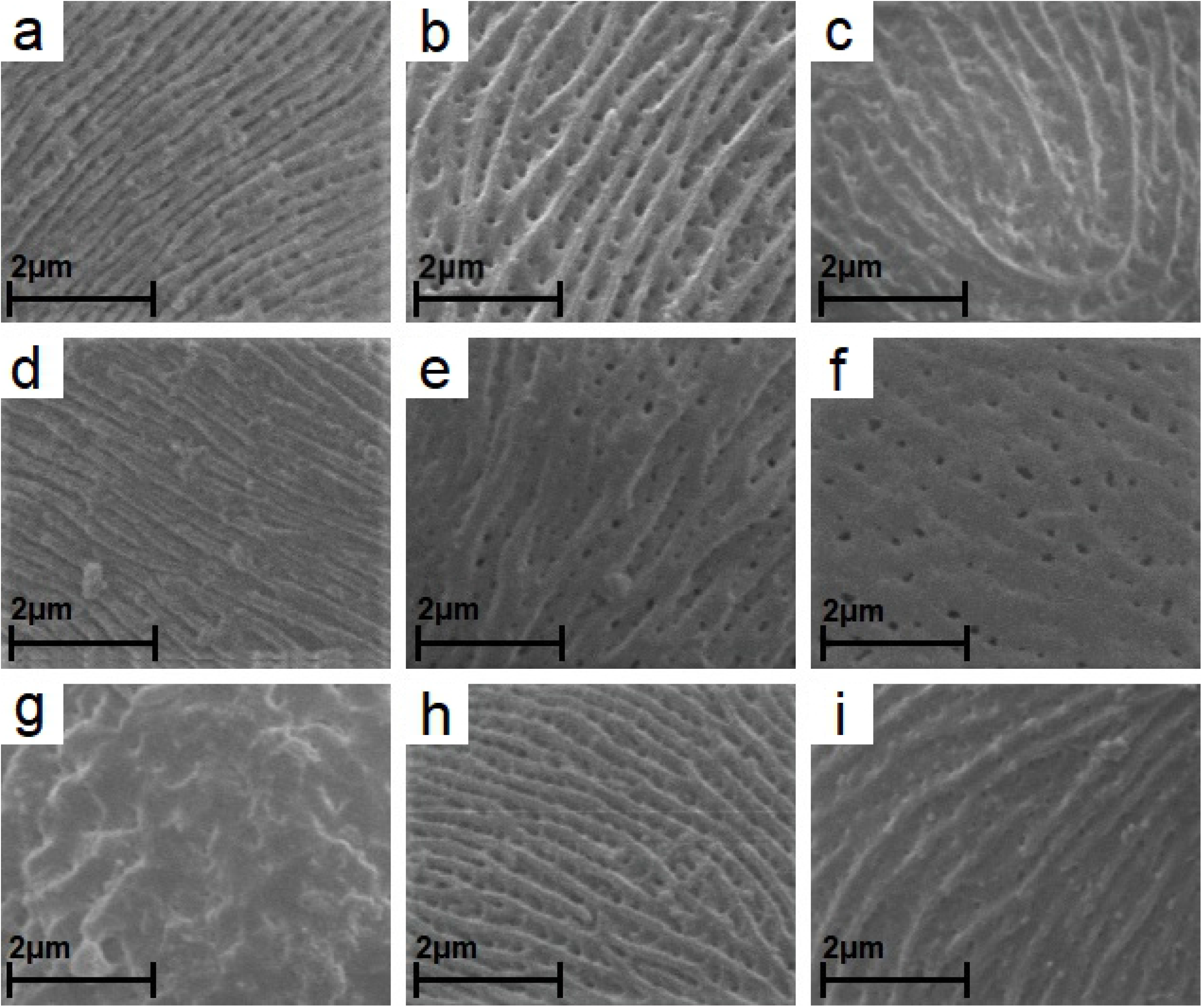
The participation of studied species in types and subtypes of striate exine ornamentation (according to Ueda [41]). (A) R. lamprocaulos (subtype – IA). (B) R. angustipaniculatus (IIA). (C) R. orthostachys (IIB). (D) R. canadensis (IIIA). (E) R. montanus (IIIB). (F) R. saxatilis (V). (G) R. odoratus (striate-verrucate ornamentation). (H) R. plicatus (IA/IIA), (I) R. apricus (IIA/IIB).

Pollen grains of the *Rubus* species studied were tricolporate, isopolar monads (Fig 1A-H). According to the pollen size classification by Erdtman [50], analysed pollen grains were medium (25.1-50 μm; 56.7%) or small (10-25 μm; 43.3%). The analysed pollen had a small range of average values for trait P, ranging from 20.57 to 30.20 μm. Therefore, most of the pollen grains belong to the upper limit of small pollen or to the lower medium-sized pollen range.

The average length of the polar axis (P) was 25.72 (18-38) μm (Fig 2, Table 3). The smallest mean P was found for pollen of *R. xanthocarpus* (20.57 μm), while the largest – for *R. dollnensis* (32.27 μm) (Fig 2, Table 3). In the *R. xanthocarpus* sample all measured pollen grains were small at a narrow range of polar axis length (18-24 μm). On the other hand, the longest pollen grains were found in *R. dollnensis* (26-38 μm).

**Table 3.**
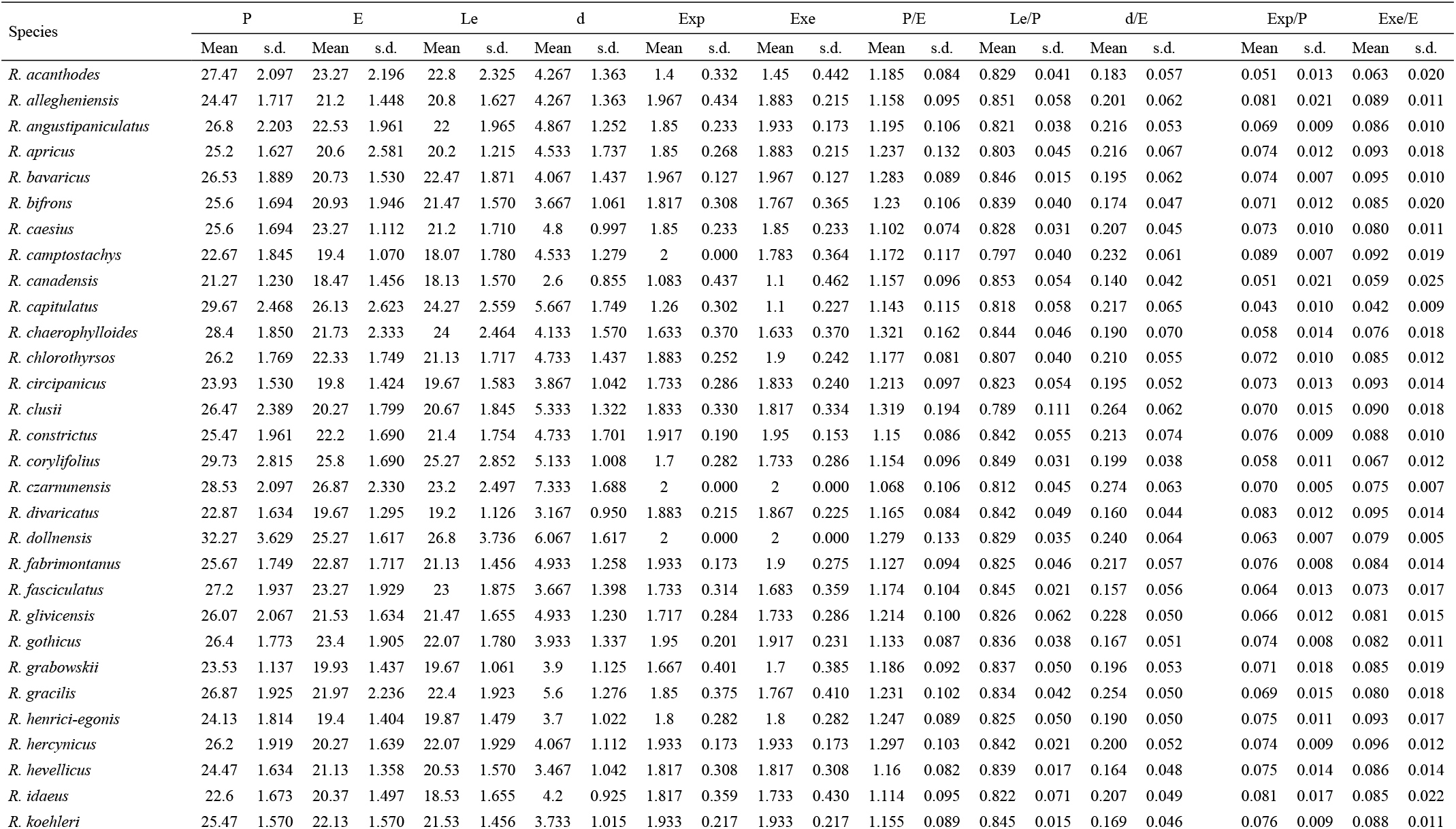
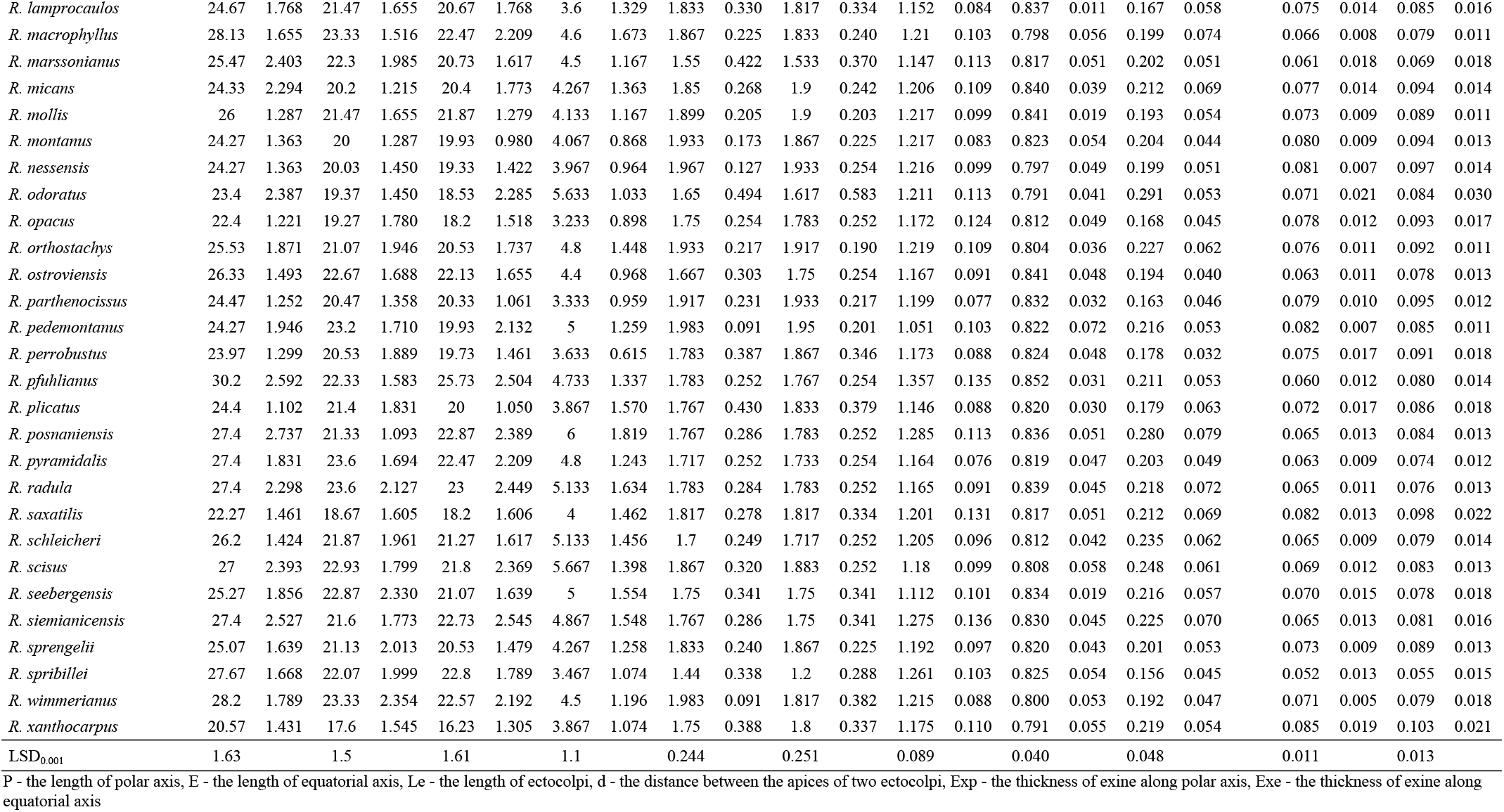
Mean values and standard deviations (s.d.) for individual species and observed traits.

| Species | P |  | E |  | Le |  | d |  | Exp |  | Exe |  | P/E |  | Le/P |  | d/E |  | Exp/P |  | Exe/E |  |
| --- | --- | --- | --- | --- | --- | --- | --- | --- | --- | --- | --- | --- | --- | --- | --- | --- | --- | --- | --- | --- | --- | --- |
|  | Mean | s.d. | Mean | s.d. | Mean | s.d. | Mean | s.d. | Mean | s.d. | Mean | s.d. | Mean | s.d. | Mean | s.d. | Mean | s.d. | Mean | s.d. | Mean | s.d. |
| *R. acanthodes* | 27.47 | 2.097 | 23.27 | 2.196 | 22.8 | 2.325 | 4.267 | 1.363 | 1.4 | 0.332 | 1.45 | 0.442 | 1.185 | 0.084 | 0.829 | 0.041 | 0.183 | 0.057 | 0.051 | 0.013 | 0.063 | 0.020 |
| *R. allegheniensis* | 24.47 | 1.717 | 21.2 | 1.448 | 20.8 | 1.627 | 4.267 | 1.363 | 1.967 | 0.434 | 1.883 | 0.215 | 1.158 | 0.095 | 0.851 | 0.058 | 0.201 | 0.062 | 0.081 | 0.021 | 0.089 | 0.011 |
| *R. angustipaniculatus* | 26.8 | 2.203 | 22.53 | 1.961 | 22 | 1.965 | 4.867 | 1.252 | 1.85 | 0.233 | 1.933 | 0.173 | 1.195 | 0.106 | 0.821 | 0.038 | 0.216 | 0.053 | 0.069 | 0.009 | 0.086 | 0.010 |
| *R. apricus* | 25.2 | 1.627 | 20.6 | 2.581 | 20.2 | 1.215 | 4.533 | 1.737 | 1.85 | 0.268 | 1.883 | 0.215 | 1.237 | 0.132 | 0.803 | 0.045 | 0.216 | 0.067 | 0.074 | 0.012 | 0.093 | 0.018 |
| *R. bavaricus* | 26.53 | 1.889 | 20.73 | 1.530 | 22.47 | 1.871 | 4.067 | 1.437 | 1.967 | 0.127 | 1.967 | 0.127 | 1.283 | 0.089 | 0.846 | 0.015 | 0.195 | 0.062 | 0.074 | 0.007 | 0.095 | 0.010 |
| *R. bifrons* | 25.6 | 1.694 | 20.93 | 1.946 | 21.47 | 1.570 | 3.667 | 1.061 | 1.817 | 0.308 | 1.767 | 0.365 | 1.23 | 0.106 | 0.839 | 0.040 | 0.174 | 0.047 | 0.071 | 0.012 | 0.085 | 0.020 |
| *R. caesius* | 25.6 | 1.694 | 23.27 | 1.112 | 21.2 | 1.710 | 4.8 | 0.997 | 1.85 | 0.233 | 1.85 | 0.233 | 1.102 | 0.074 | 0.828 | 0.031 | 0.207 | 0.045 | 0.073 | 0.010 | 0.080 | 0.011 |
| *R. camptostachys* | 22.67 | 1.845 | 19.4 | 1.070 | 18.07 | 1.780 | 4.533 | 1.279 | 2 | 0.000 | 1.783 | 0.364 | 1.172 | 0.117 | 0.797 | 0.040 | 0.232 | 0.061 | 0.089 | 0.007 | 0.092 | 0.019 |
| *R. canadensis* | 21.27 | 1.230 | 18.47 | 1.456 | 18.13 | 1.570 | 2.6 | 0.855 | 1.083 | 0.437 | 1.1 | 0.462 | 1.157 | 0.096 | 0.853 | 0.054 | 0.140 | 0.042 | 0.051 | 0.021 | 0.059 | 0.025 |
| *R. capitulatus* | 29.67 | 2.468 | 26.13 | 2.623 | 24.27 | 2.559 | 5.667 | 1.749 | 1.26 | 0.302 | 1.1 | 0.227 | 1.143 | 0.115 | 0.818 | 0.058 | 0.217 | 0.065 | 0.043 | 0.010 | 0.042 | 0.009 |
| *R. chaerophylloides* | 28.4 | 1.850 | 21.73 | 2.333 | 24 | 2.464 | 4.133 | 1.570 | 1.633 | 0.370 | 1.633 | 0.370 | 1.321 | 0.162 | 0.844 | 0.046 | 0.190 | 0.070 | 0.058 | 0.014 | 0.076 | 0.018 |
| *R. chlorothyrsos* | 26.2 | 1.769 | 22.33 | 1.749 | 21.13 | 1.717 | 4.733 | 1.437 | 1.883 | 0.252 | 1.9 | 0.242 | 1.177 | 0.081 | 0.807 | 0.040 | 0.210 | 0.055 | 0.072 | 0.010 | 0.085 | 0.012 |
| *R. circipanicus* | 23.93 | 1.530 | 19.8 | 1.424 | 19.67 | 1.583 | 3.867 | 1.042 | 1.733 | 0.286 | 1.833 | 0.240 | 1.213 | 0.097 | 0.823 | 0.054 | 0.195 | 0.052 | 0.073 | 0.013 | 0.093 | 0.014 |
| *R. clusii* | 26.47 | 2.389 | 20.27 | 1.799 | 20.67 | 1.845 | 5.333 | 1.322 | 1.833 | 0.330 | 1.817 | 0.334 | 1.319 | 0.194 | 0.789 | 0.111 | 0.264 | 0.062 | 0.070 | 0.015 | 0.090 | 0.018 |
| *R. constrictus* | 25.47 | 1.961 | 22.2 | 1.690 | 21.4 | 1.754 | 4.733 | 1.701 | 1.917 | 0.190 | 1.95 | 0.153 | 1.15 | 0.086 | 0.842 | 0.055 | 0.213 | 0.074 | 0.076 | 0.009 | 0.088 | 0.010 |
| *R. corylifolius* | 29.73 | 2.815 | 25.8 | 1.690 | 25.27 | 2.852 | 5.133 | 1.008 | 1.7 | 0.282 | 1.733 | 0.286 | 1.154 | 0.096 | 0.849 | 0.031 | 0.199 | 0.038 | 0.058 | 0.011 | 0.067 | 0.012 |
| *R. czarnunensis* | 28.53 | 2.097 | 26.87 | 2.330 | 23.2 | 2.497 | 7.333 | 1.688 | 2 | 0.000 | 2 | 0.000 | 1.068 | 0.106 | 0.812 | 0.045 | 0.274 | 0.063 | 0.070 | 0.005 | 0.075 | 0.007 |
| *R. divaricatus* | 22.87 | 1.634 | 19.67 | 1.295 | 19.2 | 1.126 | 3.167 | 0.950 | 1.883 | 0.215 | 1.867 | 0.225 | 1.165 | 0.084 | 0.842 | 0.049 | 0.160 | 0.044 | 0.083 | 0.012 | 0.095 | 0.014 |
| *R. dollnensis* | 32.27 | 3.629 | 25.27 | 1.617 | 26.8 | 3.736 | 6.067 | 1.617 | 2 | 0.000 | 2 | 0.000 | 1.279 | 0.133 | 0.829 | 0.035 | 0.240 | 0.064 | 0.063 | 0.007 | 0.079 | 0.005 |
| *R. fabrimontanus* | 25.67 | 1.749 | 22.87 | 1.717 | 21.13 | 1.456 | 4.933 | 1.258 | 1.933 | 0.173 | 1.9 | 0.275 | 1.127 | 0.094 | 0.825 | 0.046 | 0.217 | 0.057 | 0.076 | 0.008 | 0.084 | 0.014 |
| *R. fasciculatus* | 27.2 | 1.937 | 23.27 | 1.929 | 23 | 1.875 | 3.667 | 1.398 | 1.733 | 0.314 | 1.683 | 0.359 | 1.174 | 0.104 | 0.845 | 0.021 | 0.157 | 0.056 | 0.064 | 0.013 | 0.073 | 0.017 |
| *R. glivicensis* | 26.07 | 2.067 | 21.53 | 1.634 | 21.47 | 1.655 | 4.933 | 1.230 | 1.717 | 0.284 | 1.733 | 0.286 | 1.214 | 0.100 | 0.826 | 0.062 | 0.228 | 0.050 | 0.066 | 0.012 | 0.081 | 0.015 |
| *R. gothicus* | 26.4 | 1.773 | 23.4 | 1.905 | 22.07 | 1.780 | 3.933 | 1.337 | 1.95 | 0.201 | 1.917 | 0.231 | 1.133 | 0.087 | 0.836 | 0.038 | 0.167 | 0.051 | 0.074 | 0.008 | 0.082 | 0.011 |
| *R. grabowskii* | 23.53 | 1.137 | 19.93 | 1.437 | 19.67 | 1.061 | 3.9 | 1.125 | 1.667 | 0.401 | 1.7 | 0.385 | 1.186 | 0.092 | 0.837 | 0.050 | 0.196 | 0.053 | 0.071 | 0.018 | 0.085 | 0.019 |
| *R. gracilis* | 26.87 | 1.925 | 21.97 | 2.236 | 22.4 | 1.923 | 5.6 | 1.276 | 1.85 | 0.375 | 1.767 | 0.410 | 1.231 | 0.102 | 0.834 | 0.042 | 0.254 | 0.050 | 0.069 | 0.015 | 0.080 | 0.018 |
| *R. henrici-egonis* | 24.13 | 1.814 | 19.4 | 1.404 | 19.87 | 1.479 | 3.7 | 1.022 | 1.8 | 0.282 | 1.8 | 0.282 | 1.247 | 0.089 | 0.825 | 0.050 | 0.190 | 0.050 | 0.075 | 0.011 | 0.093 | 0.017 |
| *R. hercynicus* | 26.2 | 1.919 | 20.27 | 1.639 | 22.07 | 1.929 | 4.067 | 1.112 | 1.933 | 0.173 | 1.933 | 0.173 | 1.297 | 0.103 | 0.842 | 0.021 | 0.200 | 0.052 | 0.074 | 0.009 | 0.096 | 0.012 |
| *R. hevellicus* | 24.47 | 1.634 | 21.13 | 1.358 | 20.53 | 1.570 | 3.467 | 1.042 | 1.817 | 0.308 | 1.817 | 0.308 | 1.16 | 0.082 | 0.839 | 0.017 | 0.164 | 0.048 | 0.075 | 0.014 | 0.086 | 0.014 |
| *R. idaeus* | 22.6 | 1.673 | 20.37 | 1.497 | 18.53 | 1.655 | 4.2 | 0.925 | 1.817 | 0.359 | 1.733 | 0.430 | 1.114 | 0.095 | 0.822 | 0.071 | 0.207 | 0.049 | 0.081 | 0.017 | 0.085 | 0.022 |
| *R. koehleri* | 25.47 | 1.570 | 22.13 | 1.570 | 21.53 | 1.456 | 3.733 | 1.015 | 1.933 | 0.217 | 1.933 | 0.217 | 1.155 | 0.089 | 0.845 | 0.015 | 0.169 | 0.046 | 0.076 | 0.009 | 0.088 | 0.011 |
| *R. lamprocaulos* | 24.67 | 1.768 | 21.47 | 1.655 | 20.67 | 1.768 | 3.6 | 1.329 | 1.833 | 0.330 | 1.817 | 0.334 | 1.152 | 0.084 | 0.837 | 0.011 | 0.167 | 0.058 | 0.075 | 0.014 | 0.085 | 0.016 |
| *R. macrophyllus* | 28.13 | 1.655 | 23.33 | 1.516 | 22.47 | 2.209 | 4.6 | 1.673 | 1.867 | 0.225 | 1.833 | 0.240 | 1.21 | 0.103 | 0.798 | 0.056 | 0.199 | 0.074 | 0.066 | 0.008 | 0.079 | 0.011 |
| *R. marssonianus* | 25.47 | 2.403 | 22.3 | 1.985 | 20.73 | 1.617 | 4.5 | 1.167 | 1.55 | 0.422 | 1.533 | 0.370 | 1.147 | 0.113 | 0.817 | 0.051 | 0.202 | 0.051 | 0.061 | 0.018 | 0.069 | 0.018 |
| *R. micans* | 24.33 | 2.294 | 20.2 | 1.215 | 20.4 | 1.773 | 4.267 | 1.363 | 1.85 | 0.268 | 1.9 | 0.242 | 1.206 | 0.109 | 0.840 | 0.039 | 0.212 | 0.069 | 0.077 | 0.014 | 0.094 | 0.014 |
| *R. mollis* | 26 | 1.287 | 21.47 | 1.655 | 21.87 | 1.279 | 4.133 | 1.167 | 1.899 | 0.205 | 1.9 | 0.203 | 1.217 | 0.099 | 0.841 | 0.019 | 0.193 | 0.054 | 0.073 | 0.009 | 0.089 | 0.011 |
| *R. montanus* | 24.27 | 1.363 | 20 | 1.287 | 19.93 | 0.980 | 4.067 | 0.868 | 1.933 | 0.173 | 1.867 | 0.225 | 1.217 | 0.083 | 0.823 | 0.054 | 0.204 | 0.044 | 0.080 | 0.009 | 0.094 | 0.013 |
| *R. nessensis* | 24.27 | 1.363 | 20.03 | 1.450 | 19.33 | 1.422 | 3.967 | 0.964 | 1.967 | 0.127 | 1.933 | 0.254 | 1.216 | 0.099 | 0.797 | 0.049 | 0.199 | 0.051 | 0.081 | 0.007 | 0.097 | 0.014 |
| *R. odoratus* | 23.4 | 2.387 | 19.37 | 1.450 | 18.53 | 2.285 | 5.633 | 1.033 | 1.65 | 0.494 | 1.617 | 0.583 | 1.211 | 0.113 | 0.791 | 0.041 | 0.291 | 0.053 | 0.071 | 0.021 | 0.084 | 0.030 |
| *R. opacus* | 22.4 | 1.221 | 19.27 | 1.780 | 18.2 | 1.518 | 3.233 | 0.898 | 1.75 | 0.254 | 1.783 | 0.252 | 1.172 | 0.124 | 0.812 | 0.049 | 0.168 | 0.045 | 0.078 | 0.012 | 0.093 | 0.017 |
| *R. orthostachys* | 25.53 | 1.871 | 21.07 | 1.946 | 20.53 | 1.737 | 4.8 | 1.448 | 1.933 | 0.217 | 1.917 | 0.190 | 1.219 | 0.109 | 0.804 | 0.036 | 0.227 | 0.062 | 0.076 | 0.011 | 0.092 | 0.011 |
| *R. ostroviensis* | 26.33 | 1.493 | 22.67 | 1.688 | 22.13 | 1.655 | 4.4 | 0.968 | 1.667 | 0.303 | 1.75 | 0.254 | 1.167 | 0.091 | 0.841 | 0.048 | 0.194 | 0.040 | 0.063 | 0.011 | 0.078 | 0.013 |
| *R. parthenocissus* | 24.47 | 1.252 | 20.47 | 1.358 | 20.33 | 1.061 | 3.333 | 0.959 | 1.917 | 0.231 | 1.933 | 0.217 | 1.199 | 0.077 | 0.832 | 0.032 | 0.163 | 0.046 | 0.079 | 0.010 | 0.095 | 0.012 |
| *R. pedemontanus* | 24.27 | 1.946 | 23.2 | 1.710 | 19.93 | 2.132 | 5 | 1.259 | 1.983 | 0.091 | 1.95 | 0.201 | 1.051 | 0.103 | 0.822 | 0.072 | 0.216 | 0.053 | 0.082 | 0.007 | 0.085 | 0.011 |
| *R. perrobustus* | 23.97 | 1.299 | 20.53 | 1.889 | 19.73 | 1.461 | 3.633 | 0.615 | 1.783 | 0.387 | 1.867 | 0.346 | 1.173 | 0.088 | 0.824 | 0.048 | 0.178 | 0.032 | 0.075 | 0.017 | 0.091 | 0.018 |
| *R. pfuhlianus* | 30.2 | 2.592 | 22.33 | 1.583 | 25.73 | 2.504 | 4.733 | 1.337 | 1.783 | 0.252 | 1.767 | 0.254 | 1.357 | 0.135 | 0.852 | 0.031 | 0.211 | 0.053 | 0.060 | 0.012 | 0.080 | 0.014 |
| *R. plicatus* | 24.4 | 1.102 | 21.4 | 1.831 | 20 | 1.050 | 3.867 | 1.570 | 1.767 | 0.430 | 1.833 | 0.379 | 1.146 | 0.088 | 0.820 | 0.030 | 0.179 | 0.063 | 0.072 | 0.017 | 0.086 | 0.018 |
| *R. posnaniensis* | 27.4 | 2.737 | 21.33 | 1.093 | 22.87 | 2.389 | 6 | 1.819 | 1.767 | 0.286 | 1.783 | 0.252 | 1.285 | 0.113 | 0.836 | 0.051 | 0.280 | 0.079 | 0.065 | 0.013 | 0.084 | 0.013 |
| *R. pyramidalis* | 27.4 | 1.831 | 23.6 | 1.694 | 22.47 | 2.209 | 4.8 | 1.243 | 1.717 | 0.252 | 1.733 | 0.254 | 1.164 | 0.076 | 0.819 | 0.047 | 0.203 | 0.049 | 0.063 | 0.009 | 0.074 | 0.012 |
| *R. radula* | 27.4 | 2.298 | 23.6 | 2.127 | 23 | 2.449 | 5.133 | 1.634 | 1.783 | 0.284 | 1.783 | 0.252 | 1.165 | 0.091 | 0.839 | 0.045 | 0.218 | 0.072 | 0.065 | 0.011 | 0.076 | 0.013 |
| *R. saxatilis* | 22.27 | 1.461 | 18.67 | 1.605 | 18.2 | 1.606 | 4 | 1.462 | 1.817 | 0.278 | 1.817 | 0.334 | 1.201 | 0.131 | 0.817 | 0.051 | 0.212 | 0.069 | 0.082 | 0.013 | 0.098 | 0.022 |
| *R. schleicheri* | 26.2 | 1.424 | 21.87 | 1.961 | 21.27 | 1.617 | 5.133 | 1.456 | 1.7 | 0.249 | 1.717 | 0.252 | 1.205 | 0.096 | 0.812 | 0.042 | 0.235 | 0.062 | 0.065 | 0.009 | 0.079 | 0.014 |
| *R. scisus* | 27 | 2.393 | 22.93 | 1.799 | 21.8 | 2.369 | 5.667 | 1.398 | 1.867 | 0.320 | 1.883 | 0.252 | 1.18 | 0.099 | 0.808 | 0.058 | 0.248 | 0.061 | 0.069 | 0.012 | 0.083 | 0.013 |
| *R. seebergensis* | 25.27 | 1.856 | 22.87 | 2.330 | 21.07 | 1.639 | 5 | 1.554 | 1.75 | 0.341 | 1.75 | 0.341 | 1.112 | 0.101 | 0.834 | 0.019 | 0.216 | 0.057 | 0.070 | 0.015 | 0.078 | 0.018 |
| *R. siemianicensis* | 27.4 | 2.527 | 21.6 | 1.773 | 22.73 | 2.545 | 4.867 | 1.548 | 1.767 | 0.286 | 1.75 | 0.341 | 1.275 | 0.136 | 0.830 | 0.045 | 0.225 | 0.070 | 0.065 | 0.013 | 0.081 | 0.016 |
| *R. sprengelii* | 25.07 | 1.639 | 21.13 | 2.013 | 20.53 | 1.479 | 4.267 | 1.258 | 1.833 | 0.240 | 1.867 | 0.225 | 1.192 | 0.097 | 0.820 | 0.043 | 0.201 | 0.053 | 0.073 | 0.009 | 0.089 | 0.013 |
| *R. spribillei* | 27.67 | 1.668 | 22.07 | 1.999 | 22.8 | 1.789 | 3.467 | 1.074 | 1.44 | 0.338 | 1.2 | 0.288 | 1.261 | 0.103 | 0.825 | 0.054 | 0.156 | 0.045 | 0.052 | 0.013 | 0.055 | 0.015 |
| *R. wimmerianus* | 28.2 | 1.789 | 23.33 | 2.354 | 22.57 | 2.192 | 4.5 | 1.196 | 1.983 | 0.091 | 1.817 | 0.382 | 1.215 | 0.088 | 0.800 | 0.053 | 0.192 | 0.047 | 0.071 | 0.005 | 0.079 | 0.018 |
| *R. xanthocarpus* | 20.57 | 1.431 | 17.6 | 1.545 | 16.23 | 1.305 | 3.867 | 1.074 | 1.75 | 0.388 | 1.8 | 0.337 | 1.175 | 0.110 | 0.791 | 0.055 | 0.219 | 0.054 | 0.085 | 0.019 | 0.103 | 0.021 |
| LSD<sub>0.001</sub> | 1.63 |  | 1.5 |  | 1.61 |  | 1.1 |  | 0.244 |  | 0.251 |  | 0.089 |  | 0.040 |  | 0.048 |  | 0.011 |  | 0.013 |  |

P - the length of polar axis, E - the length of equatorial axis, Le - the length of ectocolpi, d - the distance between the apices of two ectocolpi, Exp - the thickness of exine along polar axis, Exe - the thickness of exine along equatorial axis

The mean length of the equatorial diameter (E) was 21.66 (14-32) μm. The shortest mean equatorial diameter was recorded in pollen of *R. canadensis* (18.47 μm), while the longest was found in *R. czarnunensis* (26.87 μm; Table 3).

The outline in the polar view was mostly circular, triangular with obtuse apices, more rarely elliptic, whereas in the equatorial view the outline was mostly elliptic, rarely circular (Fig 1).

The mean P/E ratio was 1.19, ranging from 0.85 in *R. pedemontanus* to 1.71 in *R. saxatilis* (Table 3). On average the P/E ratio values were always above 1 and they ranged from 1.05 in *R. pedemontanus* to 1.32 in *R. chaerophylloides.* Pollen grains of the species examined were most frequently subprolate (57.3% – 997 pollen grains) or prolate-spheroidal (24.3% – 422), rarely prolate (8.9% – 155) or spheroidal (8.6% – 150) and very rarely oblate-spheroidal (0.7% – 12) and perprolate (0.2% – 4). The highest number of subprolate pollen grains was recorded in *R. henriciegonis* and *R. montanus* (each at 80%, – 24 grains), of prolate-spheroidal pollen – in *R. idaeus* (53.3% – 16 grains) and of prolate grains – in *R. chaerophylloides* (50% – 15).

The exine was two-layered, with the ectexine and endexine of about the same thickness. Mean exine thickness was 1.79 (0.5-4.0) μm; on average Exp – 1.79 μm and Exe – 1.78 μm. The exine was the thinnest in *R. canadensis* (Exp – 0.8 μm; Exe – 1.1 μm), while it was the thickest in *R. czarnuensis* and *R. dollensis* (Exp and Exe – 2.0 μm; Table 3). The relative thickness of the exine (Exp/P ratio) averaged 0.07 (0.02-0.18) and (Exe/E ratio) 0.08 (0.02-0.14). The above results were similar, indicating a more or less equal exine thickness along the entire pollen grain (Table 3).

In all the studied species, exine ornamentation was striate-perforate and very rarely striate, with the exception of *R. odoratus*, which had a striate-verrucate ornamentation with small perforations (Fig 3). Exine ornamentation elements were highly variable (Fig 3). Striae and grooves usually ran parallel to colpori and the polar axis, but frequently they also formed fingerprint-like twists. Striae were straight or forked and of varying length, width and height.

The investigated pollen of the individual *Rubus* species was classified according to the striate exine ornamentation classification proposed by Ueda [41] into four types (I-III and V) and five subtypes (I A, II A,B and III A,B). The cited author distinguished six types (I-VI) and six subtypes (I-III, each A and B). In our study types IV, VI and subtype IB were not found (Fig 3, Table 4). The greatest number of species (18) belonged to the IIA subtype, which was characterised by fairly distinct striae, narrow grooves and frequently by prominent, numerous perforations. Subtypes IA, IIA/IIB, IIB and IIIA were represented by a relatively large number of species (8, 11, 8 and 9 species, respectively), while types IA/IIA, IIIB and V – by only one species. Among the 58 examined species, 12 had two types of exine ornamentation (Fig 3, Table 4).

**Table 4.** Striate exine ornamentation types and subtypes of studied Rubus species (according to Ueda [41] classification).

| Striate exine ornamentation type or subtype | Species |
| --- | --- |
| IA | *R. chaerophylloides*, *R. corylifolius*, *R. fasciculatus*, *R. henrici-egonis*, *R. hercynicus*, *R. lamprocaulos*, *R. pfuhlianus*, *R. posnaniensis* |
| IA/IIA | *R. plicatus* |
| IIA | *R. acanthodes*, *R. allegheniensis*, *R. angustipaniculatus*, *R. camptostachys*, *R. circipanicus*, *R. constrictus*, *R. grabowskii*, *R. gracilis*, *R. hevellicus*, *R. koehleri*, *R. macrophyllus*, *R. marssonianus*, *R. nessensis*, *R. ostroviensis*, *R. parthenocissus*, *R. sprengelii*, *R. wimmerianus*, *R. xanthocarpus* |
| IIA/IIB | *R. apricus*, *R. bavaricus*, *R. bifrons*, *R. capitulatus*, *R. clusii*, *R. micans*, *R. pyramidalis*, *R. spribillei*, *R. chlorothyrsos*, *R. schleicheri*, *R. seebergensis* |
| IIB | *R. caesius*, *R. dollnensis*, *R. glivicensis*, *R. gothicus*, *R. idaeus*, *R. mollis*, *R. orthostachys*, *R. siemianicensis* |
| IIIA | *R. canadensis*, *R. czarnunensis*, *R. divaricatus*, *R. fabrimontanus*, *R. opacus*, *R. pedemontanus*, *R. perrobustus*, *R. radula*, *R. scissus* |
| IIIB | *R. montanus* |
| striate-verrucate | *R. odoratus* |
| V | *R. saxatilis* |

In most of the species (56 of the 58), elliptic or circular perforations of different diameters (0.05-0.4 μm) were found at the bottom of the grooves (Fig 3). The perforations were not found in *R. canadensis* and *R. czarnunensis.* In the majority of the species studied the perforations were small, with similar diameters (0.1-0.2 μm) and more or less numerous, with the exception of *R. bifrons, R. capitulatus, R. constrictus, R. gracilis, R. hercynicus, R. lamprocaulos, R. odoratus, R. opacus, R. orthostachys, R. ostroviensis, R. pedemontanus, R. perrobustus* and *R. radula*, where they were relatively few. The single perforations were observed in *R. corylifolius, R. czarnunensis, R. henrici-egonis* and *R. pyramidalis.*

Pollen grains usually had three apertures – colpori. Ectoapertures – colpi were arranged meridionally, regularly, they were more or less evenly spaced and long, at a mean length of 21.23 (14-32) μm (Table 3; Fig 4D-F). On average, the length of colpi constituted 83% (from 60 to 100%) of the polar axis length, with the shortest colpi found in *R. xanthocarpus* (16.2 μm) and the longest in *R. corylifolius* (25.3 μm). Colpi were fusiform in outline. Their width was variable and usually greatest in the equatorial region. Sculpturing of ectocolpus membranes approached rugulate, rarely partly psilate (Fig 4D-F). Colpus margins frequently had small undulations (Fig 4D-F).

**Fig 4.**
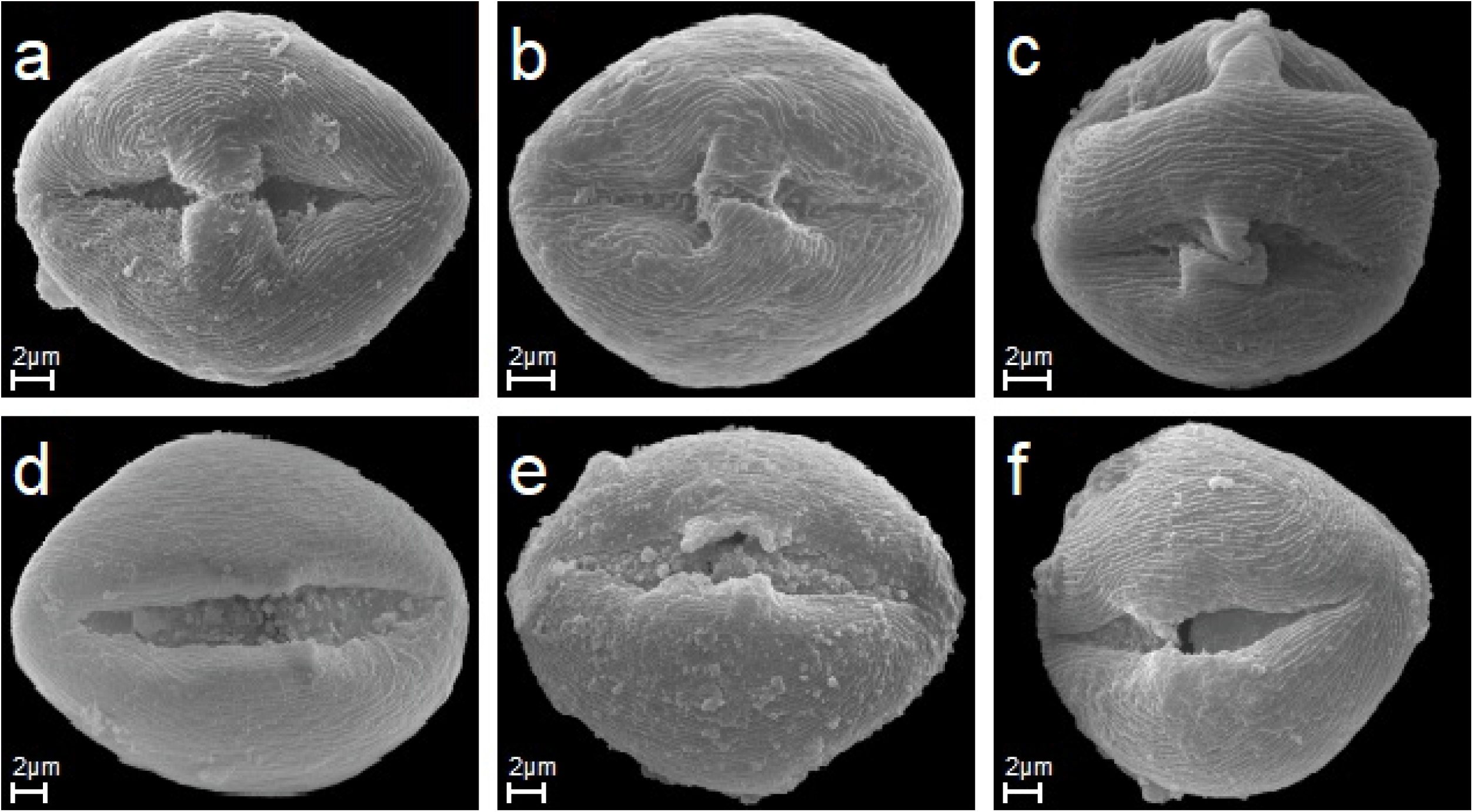
The bridge and apertures of studied species. A-C. R. macrophyllus, R. circipanicus, R. angustipaniculatus the bridge (exine connection between the margins of an aperture – colporus) in three pollen grains in equatorial view. D-F. R. gothicus, R. scisus, R. nessensis colporus with rugulate membrane in three pollen in equatorial view.

In all of the species studied the colpus was crossed at the equator by a bridge dividing it into two parts, formed by two bulges of the ectexine that meet in the middle (Fig 4A-C). The bulges were of the same or unequal length.

The polar area index (PAI) or the apocolpium index (d/E ratio) averaged 0.20 (0.08-0.45). The lowest mean values of this index were recorded in *R. canadensis* (0.14), while the highest – in *R. odoratus* (0.29) (Table 3).

Endoapertures were usually located in the middle of colpi, less frequently asymmetrically, usually singly and very rarely in pairs. They were circular or elliptic in outline with irregular margins (Fig 4D-F).

### Pollen key

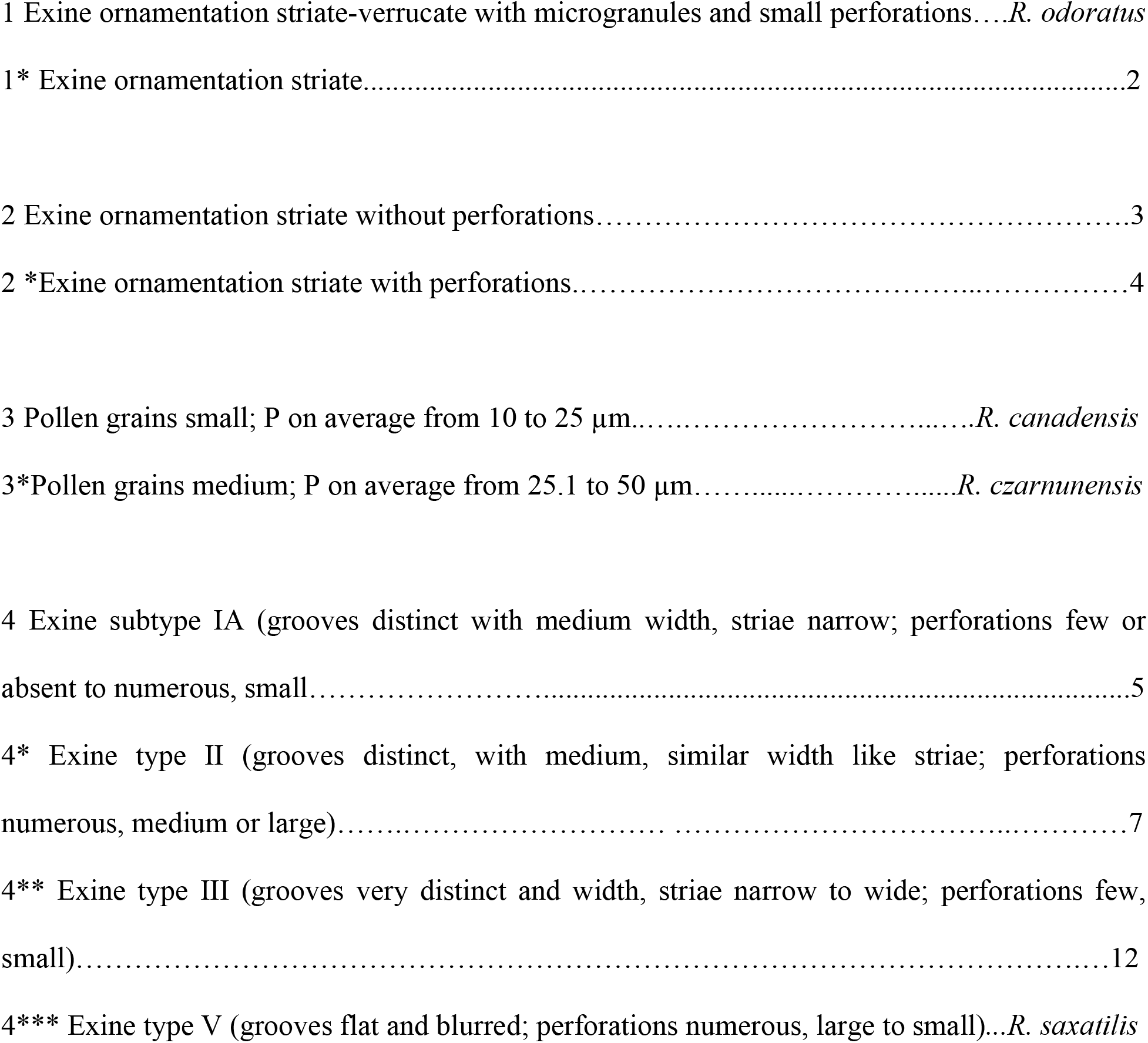
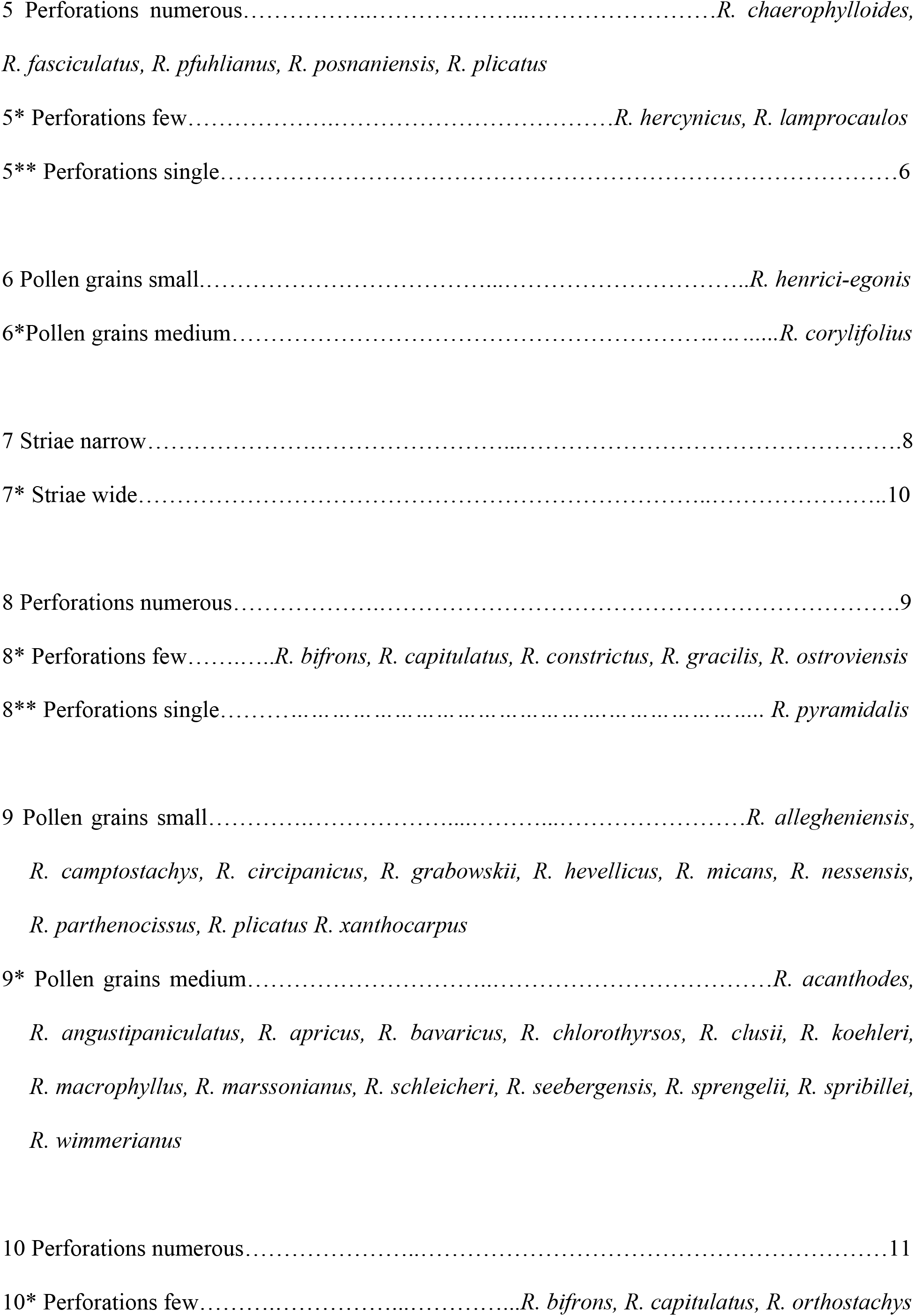
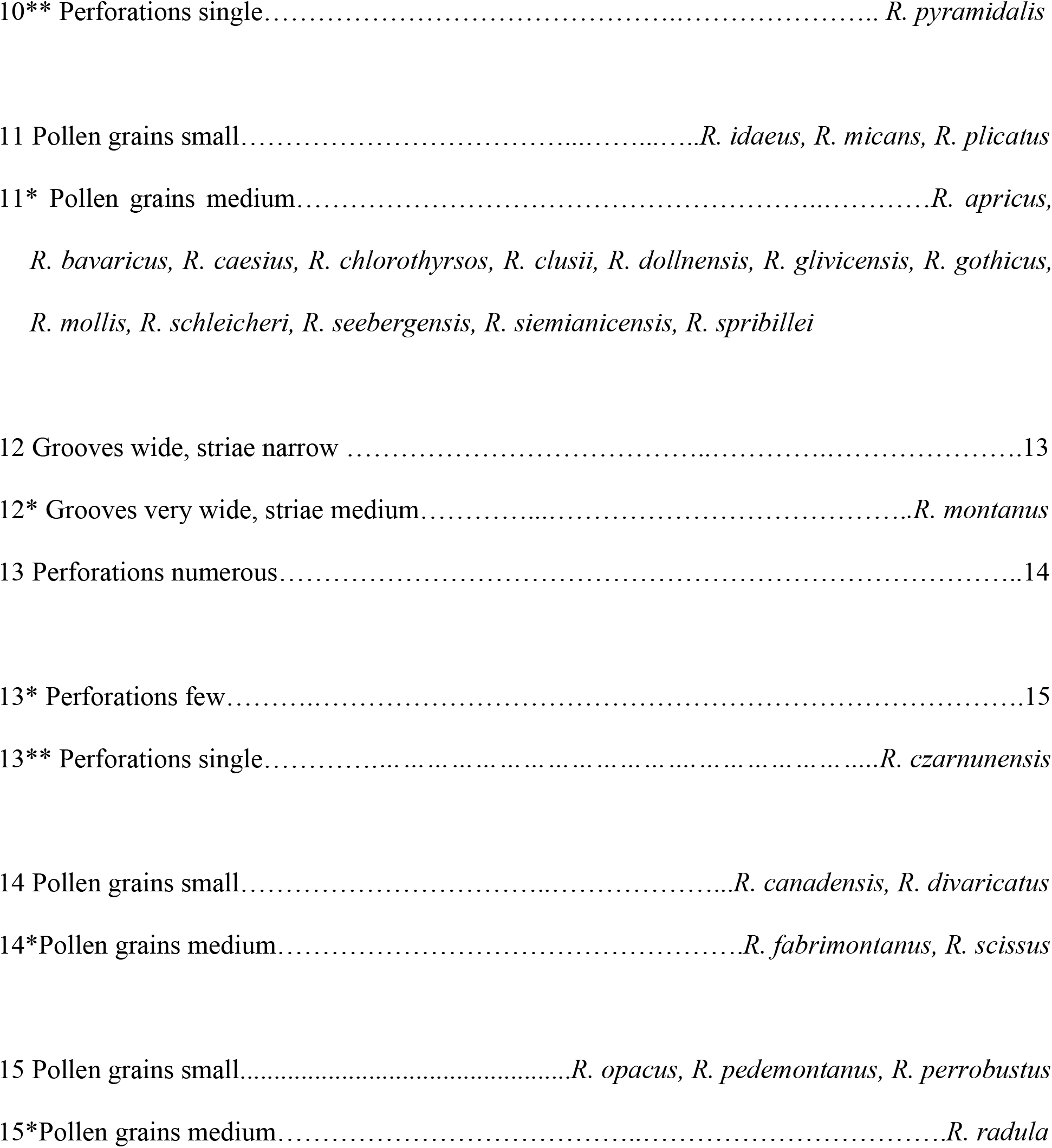

### Intrageneric and interspecific variability of pollen grains

The results of the MANOVA indicated that all the species were significantly different with regard to all of the 11 quantitative traits (Wilk’s *λ*=0.04048; *F*_627,18111_=9.98; *P*<0.0001). The analysis of variance for the 11 biometric traits [P (*F*_57,1682_=40.42), E (*F*_57,1682_=33.51), Le (*F*_57,1682_=32.48), d (*F*_57,1682_=12.41), Exp (*F*_57,1682_=11.26), Exe (*F*_57,1682_=12.11), P/E (*F*_57,1682_=9.87), Le/P (*F*_57,1682_=3.89) d/E (*F*_57,1682_=9.24), Exp/P (*F*_57,1682_=15.35) and Exe/E (*F*_57,1682_=15.29)] confirmed variability of the tested species at a significance level α=0.001. The mean values and standard deviations for the observed traits indicated a high variability among the tested species, for which significant differences were found in terms of all the analysed morphological traits (Table 3).

The correlation analysis indicated statistically significant correlation coefficients for 25 out of 55 coefficients (Table 5). A total of 16 out of 25 significantly correlated pairs of traits were characterised by positive correlation coefficients. In the case of 30 pairs of traits, no significant correlation was established.

**Table 5.** Correlation coefficients between all pairs of observed traits.

| Trait | P | E | Le | d | Exp | Exe | P/E | Le/P | d/E | Exp/P | Exe/E |
| --- | --- | --- | --- | --- | --- | --- | --- | --- | --- | --- | --- |
| P | 1 |  |  |  |  |  |  |  |  |  |  |
| E | 0.820*** | 1 |  |  |  |  |  |  |  |  |  |
| Le | 0.975*** | 0.799*** | 1 |  |  |  |  |  |  |  |  |
| d | 0.575*** | 0.614*** | 0.477*** | 1 |  |  |  |  |  |  |  |
| Exp | 0.015 | 0.015 | -0.014 | 0.186 | 1 |  |  |  |  |  |  |
| Exe | -0.034 | -0.028 | -0.045 | 0.156 | 0.937*** | 1 |  |  |  |  |  |
| P/E | 0.322* | -0.275* | 0.310* | -0.026 | 0 | -0.012 | 1 |  |  |  |  |
| Le/P | 0.169 | 0.141 | 0.380** | -0.285* | -0.139 | -0.075 | 0.028 | 1 |  |  |  |
| d/E | 0.238 | 0.17 | 0.124 | 0.878*** | 0.226 | 0.207 | 0.143 | -0.454*** | 1 |  |  |
| Exp/P | -0.632*** | -0.520*** | -0.641*** | -0.22 | 0.757*** | 0.730*** | -0.201 | -0.236 | 0.033 | 1 |  |
| Exe/E | -0.533*** | -0.635*** | -0.537*** | -0.245 | 0.710*** | 0.779*** | 0.157 | -0.184 | 0.07 | 0.892*** | 1 |

\* P<0.05; \*\* P<0.01; \*\*\* P<0.001
P - the length of polar axis, E - the length of equatorial axis, Le - the length of ectocolpi, d - the distance between the apices of two ectocolpi, Exp - the thickness of exine along polar axis, Exe - the thickness of exine along equatorial axis

In the presented dendrogram, as a result of agglomeration grouping using the Euclidean distance method, all the examined *Rubus* species were divided into four groups (Fig 5). The first group (I) comprised one species – *R. czarnunensis*, while the second one (II) four species (*R. dollnensis, R. corylifolius, R. chaerophylloides* and *R. phuhianus*). The third group was divided into two subgroups: III A – *R. camptostachys, R. xanthocarpus, R. clussi, R. odoratus*, and III B – including all the other species from this group. The fourth group (IV) comprised *R. canadensis, R. capitulatus, R. acanthoides* and *R. spribillei.*

**Fig 5.**
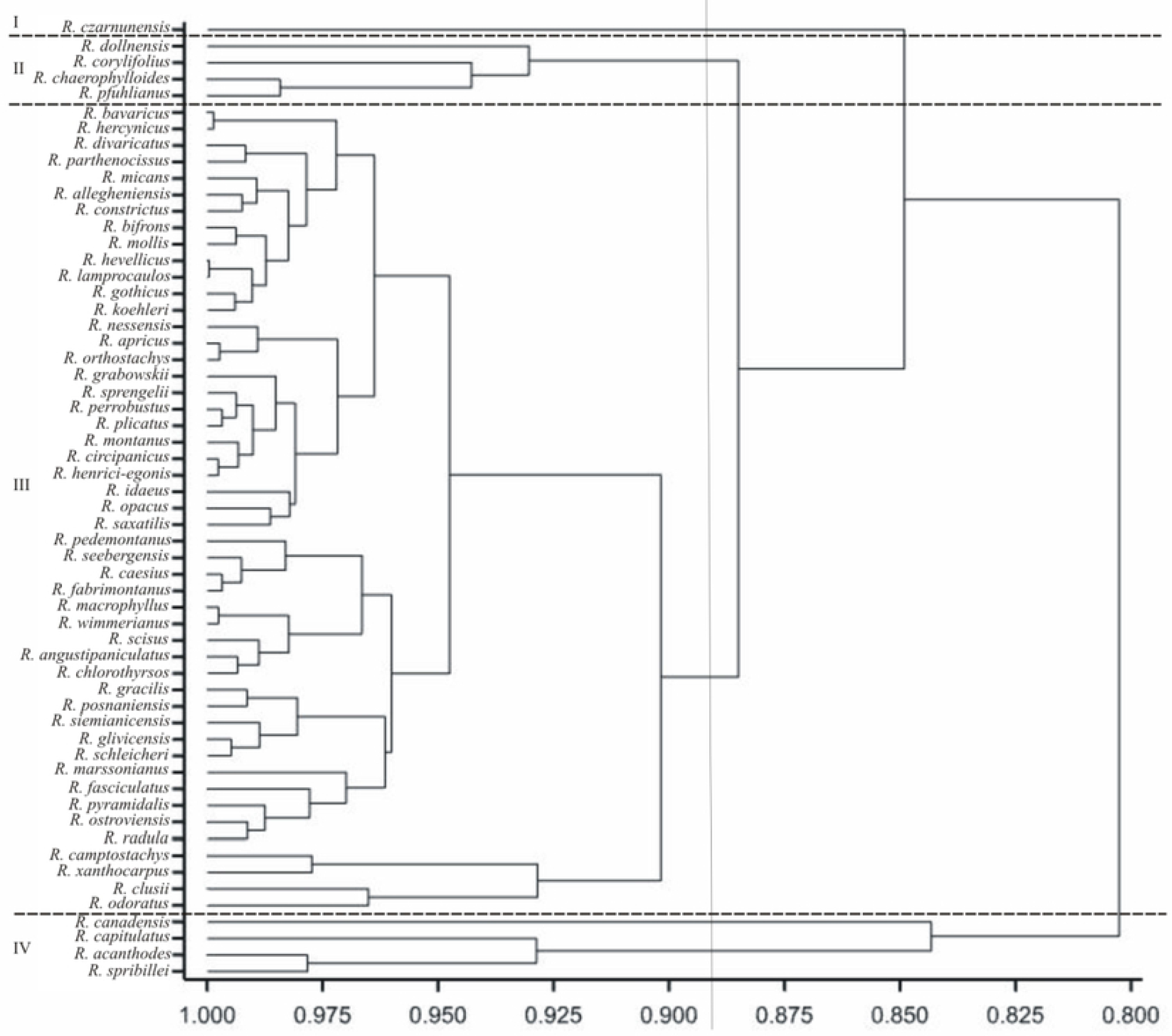
Dendrogram of cluster groupings of Rubus species based on all 11 morphological traits.

Individual traits were of varying importance and had different shares in the joint multivariate variation. A study on the multivariate variation for species includes also identification of the most important traits in the multivariate variation of species. Analysis of canonical variables is a statistical tool making it possible to solve the problem of multivariate relationships. Figure 6 shows the variability of the pollen grain features in 58 studied *Rubus* species in terms of the first two canonical variables. In the graph the coordinates of the point for particular shrubs were the values for the first and second canonical variable, respectively. The first two canonical variables accounted for 56.75% of the total multivariate variability between the individual species. Five groups of species were distinguished (Fig 5). A majority of the examined species were found in the first group (I), which means that they had more or less similar pollen features. Only one up to maximum three species (II – *R. capitulatus*, III – *R. xantocarpus*, IV – *R. acanthoides* and *R. spribillei*, and V – *R. corylifolius, R. dollnensis*, and *R. czarnunensis*) fell into the other four groups (Fig 6). Pollen grains of *R. capitulatus* were the most different from those of the other species (large, with a thin exine and the P/E ratio usually prolate-spheroidal). Species from groups IV and V had the largest pollen grains and *R. xantocarpus* (group III) – the smallest ones.

**Fig 6.**
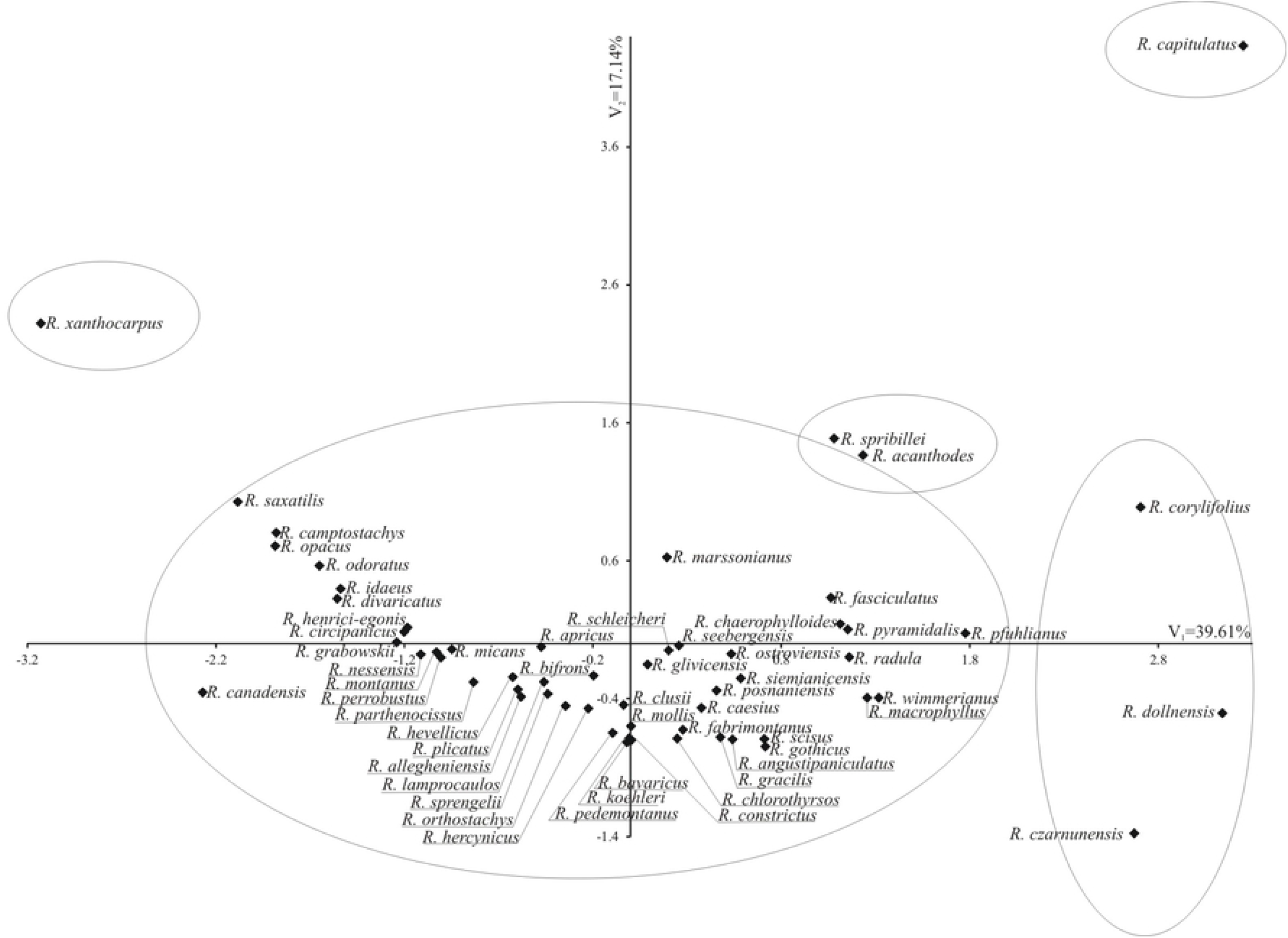
Distribution of the studied Rubus species in the space of the first two canonical variables.

The most significant, positive, linear relationship between the first canonical variables was found for P, E, Le and d, while it was negative for Exp/P and Exe/E (Table 6). The second canonical variable was significantly negatively correlated with Exp, Exe, Exp/P and Exe/E (Table 6). The greatest variation in terms of all the traits jointly (measured Mahalanobis distances) was found for *R. canadensis* and *R. capitulates* (the Mahalanobis distance between them amounted to 8.24). The greatest similarity was found for *R. lamprocaulos* and *R. hevellicus* (0.313).

**Table 6.** Correlation coefficients between the first two canonical variables and original traits.

| Trait | First canonical variable | Second canonical variable |
| --- | --- | --- |
| P | 0.9634*** | -0.0536 |
| E | 0.9353*** | -0.0382 |
| Le | 0.9427*** | -0.0812 |
| d | 0.5995*** | -0.1054 |
| Exp | -0.0477 | -0.5907*** |
| Exe | -0.0993 | -0.6587*** |
| P/E | 0.0751 | -0.0254 |
| Le/P | 0.1822 | -0.1743 |
| d/E | 0.1939 | -0.087 |
| Exp/P | -0.6568*** | -0.3354* |
| Exe/E | -0.6497*** | -0.3919** |
| Percentage of explained multivariate variability | 39.61% | 17.14% |

\* P<0.05; \*\* P<0.01; \*\*\* P<0.001
P - the length of polar axis, E - the length of equatorial axis, Le - the length of ectocolpi, d - the distance between the apices of two ectocolpi, Exp - the thickness of exine along polar axis, Exe - the thickness of exine along equatorial axis

## Discussion

Similarly to a majority of palynologists, the authors of this study maintain that exine ornamentation features were diagnostic for pollen grains of the *Rubus* species [25–28, 30–32, 35, 39, 40, 42, 43, 58, 59]. The most important exine ornamentation traits include the width, number and course of grooves (muri) and the width of the striae as well as the number and diameter of perforations [31, 32, 35, 42, 43, 59–61]. Some authors considered pollen size and shape as potentially important features in the diagnosis of the analysed genus [30, 32, 58], while others claim that they have no diagnostic significance [31, 42, 44]. Based on our results, we partially agree with the opinion of these former, because the length of the polar axis (P) has been an important feature.

In a study by Li et al. [43] the 103 examined *Rubus* species from China belonged to four types of exine ornamentation (rugulate, striate, cerebroid and reticulate-perforate), which were further divided into 11 subtypes. Other palynologists distinguish in brambles mainly striate or striate-perforate exine ornamentation [25, 27, 28, 30–32, 34, 35, 39, 40, 42, 59]. Except for the typical striate ornamentation, also striate-scabrate, striate-rugulate or rugulate [31, 42], echinate or gemmate [25], verrucate [25, 39, 40], baculate and clavate [27, 28] or reticulate ornamentation [59] have been rarely observed. According to current palynological studies, European bramble species are slightly less variable in terms of this feature than Asian ones. Our results confirm this thesis, because in the examined pollen grains only two types of exine ornamentation (striate and striate-verrucate with microgranules) were found.

Ueda & Tomita [60] and Ueda [41] distinguished six types and six subtypes of exine ornamentation in species and other taxa from the genus *Rosa* and the family Rosaceae, including the genus *Rubus*. In the current study they were classified into four types (types IV and VI were not identified) and five subtypes (I A, II A, B, III A, B). Our results were similar to the cited authors, since most of the examined pollen belonged to the IIA and IIIA subtypes and no grains were found in the very rarely represented types IV and VI or subtype IB. The only species described both by Ueda [41] and in our study was *R. odoratus*. Ueda [41] described it as a type VI and we as type V.

The research results obtained in this study confirmed the diagnostic significance of the number and diameter of perforations, found by Hebda & Chinnappa [39, 40], Monasterio-Huelin & Pardo [30], Tomlik-Wyremblewska [31], Li et al. [43], Wrońska-Pilarek et al. [32] or Ghosh & Saha [59], because these traits allowed to distinguish certain *Rubus* species (see: pollen key). On the other hand, groups of species from different sections possess similar numbers of perforations (e.g. *R. opacus* from the series *Rubus*, *R. canadensis* from the series *Canadenses* or *R. henrici-egonis* from the series *Discolores*). However, also species from many different sections (e.g. *Rubus, Alleghenienses, Sylvatici* or *Micantes*) representing the subgenus *Rubus* were characterised by high numbers of small perforations with similar diameters. Hebda and Chinnappa [39] distinguished two types of perforations in the family Rosaceae (striate – macroperforate and non-striate – macroperforate, each with six subtypes) possibly indicating different evolutionary lines. According to the above cited study, pollen of *Rosa* (with *Prunus*, *Rubus* and *Spiraea*) belongs to the subcategory with striae separated by grooves, containing larger perforations (0.1-0.2 μm in diameter). The current data corroborated this latter thesis, with the reservation that some of the species were characterised by ornamentation different than striate (*R. odoratus* – striate-verrucate with microgranules), and that perforation diameters in *Rubus* ranged from 0.05 to 0.4 μm. In turn, Hebda and Chinnappa [40] classified pollen types in Rosaceae into six main categories: 1 – striate and macroperforate, 2 – striate and microperforate, 3 – tuberculate and perforate, 4 – microverrucate, 5 – verrucate and 6 – perforate, without supratectal features. They included species from the *Rubus* genus, similarly to the study from 1990, in type 1 (striae long and parallel to colpus). Our studies demonstrated that the inclusion of the *Rubus* genus into one type is too general because, firstly, there were bramble species with the striate-verrucate exine ornamentation with microgranules (e.g. *R. odoratus*), with perforations sometimes being large, but also small (type 2 – striate and microperforate). Additionally, in some species perforations were very scarce or did not occur at all (e.g. *R. corylifolius, R. henriciegonis, R. canadensis, R. czarnuensis*). Consequently, species from the *Rubus* genus also belong to other types mentioned above, as well as types not mentioned by Hebda & Chinnappa [40].

In the opinion of many researchers the bridge was present in most of the studied *Rubus* species [30–32, 42]. They were wide, well-developed and with margins. In brambles Tomlik-Wyremblewska [31] distinguished two bridge types, with margins stretched or constricted at the equator. In our study, bridges were observed in all the analysed bramble species and this structure was not used as a basis for the identification of species, because its characteristics were too similar. Besides, it usually appeared in mature pollen grains, so it could not be noticed when analysing pollen at other developmental stages.

The presented results shows that out of 1740 measured pollen grains, 753 (43.3%) were small (15-25 μm) and 987 (56.7%) were medium (25-50 μm). Although on average 38 species had medium-sized pollen and 20 species – small-sized ones, the average length of the polar axis (P) rarely exceeded the limit medium value of 25 μm. Similar results were obtained by most other researchers [27, 28, 30–32, 35, 42, 43, 58, 59].

In the opinion of Li et al. [43] pollen shape varied from spheroidal, subspheroidal, prolate and perpolate, to occasionally rhomboid and hexagonal. In turn, Monasterio-Huelin & Pardo [30] stated that they were just prolate or spheroidal, while other authors distinguished several pollen shape types – subprolate, prolate spheroidal, prolate or perprolate [31, 32, 34, 35, 42, 59]. We agree with Tomlik-Wyremblewska [31, 42] that pollen shape turned out to be a poor criterion in identifying bramble species, because most pollen grains (81.6%) have a similar shape – subprolate or prolate-spheroidal.

The arrangement of the investigated species on the dendrogram (Fig 5) does not corroborate the division of the genus *Rubus* into subgenera, sections and series [16], currently adopted in taxonomy. Species from three different subgenera (*R. saxatilis* and *R. xanthocarpus* from the subgenus *Cylactis*, *R. odoratus* from the subgenus *Anoplobatus* and *R. idaeus* from the subgenus *Idaeobatus*) were found in the same group III, with most of the species from a large subgenus *Rubus*. Similar results were obtained for the three sections from the subgenus *Rubus* (*Rubus, Corylifolii* and *Caesii*). Thus, *R. caesius* from the section *Caesii* and *R. gothicus, R. camptostachys, R. mollis* or *R. fabrimontanus* from the section *Corylifolii* were found in group III, with the species representing the most numerous third section of *Rubus*. Also in the case of the series it were not observed that species belonging to these taxa formed separate groups (Figs 5, 6). Other genera of the family Rosaceae (e.g. *Spiraea, Rosa, Crataegus*) showed a correlation between pollen morphology and intrageneric taxonomic classification [62, 63, 64]. In *Rubus* the lack of dependence could be the result of apomixis, defined as the replacement of the normal sexual reproduction by asexual reproduction, without fertilisation, which could reduce natural variability.

## Acknowledgements

We kindly thank Nuala Scanlon-Mederski (an English native proofreader) for linguistic support.

## References

1. Gustafsson A. The genesis of the European blackberry flora. Acta Univ Lund. 1943; 39: 1–200.

2. Kurtto A, Weber HE, Lampinen R, Sennikov AN. Atlas Florae Europaeae: Distribution of vascular plants in Europe: Rosaceae (Rubus). Helskinki: The Committee for Mapping the Flora of Europe & Societas Biologica Fennica Vanamo; 2010.

3. Focke WO. Synopsis Ruborum Germaniae: Die deutschen Brombeerarten ausführlich beschrieben und erläutert. Bremen: C. Ed. Müllers’s Verlagsbuchhandlung; 1877.

4. Focke WO. Rosaceae. In: Engler A, Prantl K, editors. Die Natürlichen Pflanzenfamilien III, Leipzig: W. Engelmann; 1894.

5. Robertson KR. The genera of Rosaceae in the southeastern United States. J Arnold Arbor. 1974; 55: 352–360.

6. Jennings DL. Raspberries and Blackberries. Their Breeding, Diseases and Growth. London: Academic Press; 1988.

7. Gu Y, Zhao CM, Jin W, Li WL. Evaluation of *Rubus* germplasm resources in China. Acta Hortic. 1993; 352: 317–324.

8. Thompso MM. Chromosome numbers of *Rubus* species at the National Clonal Germplasm Repository. HortScience 1995; 30: 1447–1452.

9. Weber HE. *Rubus* L. In: Hegi G, Weber HE, editors. Illustrierte Flora von Mitteleuropa IV/2a. 3rd edn. Berlin: Blackwell Wissenschafts-Verlag; 1995. pp. 284–595.

10. Stevens PF. 2001 onwards. Angiosperm Phylogeny Website. July 2017. Available from: http://www.mobot.org/MOBOT/research/APweb/ Cited 16 July 2019.

11. Potter D, Eriksson T, Evans RC, Oh S, Smedmark JEE, Morgan DR, et al. Phylogeny and classification of Rosaceae. Pl Syst Evol. 2007; 266: 5–43. https://doi.org/10.1007/s00606-007-0539-9

12. APG IV [Angiosperm Phylogeny Group IV]. An update of the angiosperm phylogeny group classification for the orders and families of flowering plants. Bot J Linn Soc. 2016; 181: 1–20. https://doi.org/10.1111/boj.12385

13. Focke WO. Species Ruborum monographiae generis Rubi prodromus. Bibl Bot. 1910; 17: 1–120.

14. Focke, WO. Species Ruborum monographiae generis Rubi prodromus. Bibl Bot. 1914; 17: 1–274.

15. Alice LA, Campbell ChS. Phylogeny of *Rubus* (Rosaceae) based on nuclear ribosomal DNA internal transcribed spacer region sequences. Am J Bot. 1999; 86: 81–97.

16. Zieliński J. The genus *Rubus* (Rosaceae) in Poland. Pol Bot Stud. 2004; 16: 1–300.

17. Stace CA. Plant Taxonomy and Biosystematics. 2nd ed. Cambridge: Cambridge University Press; 1989.

18. Kosiński P, Maliński T, Sliwinska E, Zieliński J. *Rubus prissanicus* (Rosaceae), a new bramble species from North West Poland. Phytotaxa 2018; 344: 239–247. http://dx.doi.org/10.11646/phytotaxa.344.3.4

19. Piwowarski B. Brambles of the Jędrzejów Plateau (Nida Basin) in the Małopolska Upland. The Polish Dendrology Society Yearbook 2013; 61: 21–27.

20. Sudre H. Rubi Europae. Paris: Librairie des Sciences Naturelles; 1917.

21. Schori M, Furness CA. Pollen diversity in Aquifoliales. Bot J Linn Soc. 2014; 175: 169–190. https://doi.org/10.1111/boj.12163

22. Song JH, Oak MK, Roh HS, Hong SP. Morphology of pollen and orbicules in the tribe Spiraeeae (Rosaceae) and its systematic implications. Grana 2017; 56: 351–367. https://doi.org/10.1080/00173134.2016.1274334

23. Almeida GS, Mezzonato-Pires AC, Mendonça CBF, Gonçalves-Esteves V. Pollen morphology of selected species of Piriqueta Aubl (Passifloraceae sensu lato). Palynology 2018; 43: 43–52. https://doi.org/10.1080/01916122.2018.1434252

24. Erdtman G, Berglund B, Praglowski J. An Introduction to a Scandinavian Pollen Flora. Grana 1961; 2: 3–86.

25. Reitsma TJ. Pollen morphology of some European Rosaceae. Acta Bot Neerl. 1966; 15: 290–379.

26. Teppner H. Zur Kenntnis der Gattung Waldsteinia L. – Schlüssel zum Bestimmen von Rosaceen Polleeinschliesslich ählicher Pollen – formen aus andere Familien. Phyton 1966; 3-4: 224–238.

27. Eide F. Key for Northwest European Rosaceae pollen. Grana 1981a; 20: 101–118.

28. Eide F. On the pollen morphology of *Rubus chamaemorus* L. (Rosaceae). Grana 1981b; 20: 25–27.

29. Gonzalez Romano ML, Candau PA. Contribution to palynological studies in the Rosaceae. Acta Bot Malac. 1989; 14: 105–116.

30. Monasterio-Huelin E, Pardo C. Pollen morphology and wall stratification in *Rubus* L. (Rosaceae) in the Iberian Peninsula. Grana 1995; 34: 229–236.

31. Tomlik-Wyremblewska A. Pollen morphology of genus *Rubus* L. Part I. Introductory studies of the European representatives of the subgenus *Rubus* L. Acta Soc Bot Pol Pol. 1995; 64: 187–203.

32. Wrońska-Pilarek D, Jagodziński AM, Maliński T. Morphological studies of pollen grains of the Polish endemic species of the genus *Rubus* L. (Rosaceae). Biologia 2012; 67: 87–96. https://doi.org/10.2478/s11756-011-0141-z

33. Wrońska-Pilarek D, Danielewicz W, Bocianowski J, Maliński T, Janyszek M. Comparative Pollen Morphological Analysis and Its Systematic Implications on Three European Oak (*Quercus* L., Fagaceae) Species and Their Spontaneous Hybrids. PLoS One. 2016; 11: 1–19. https://doi.org/10.1371/journal.pone.0161762

34. Kasalkheh R, Jorjani E, Sabouri H, Habibi M, Sattarian A. Pollen morphology of the genus *Rubus* L. subgenus Rubus (Rosaceae) in Iran. Nova Bioliogica Reperta 2017; 4: 9–18.

35. Wrońska-Pilarek D, Maliński T, Lira J. 2006. Pollen morphology of Polish species of genus *Rubus* L. -*Rubus gracilis* J. Presl & C Presl Dendrobiology. 2006; 56: 69–77.

36. Naruhashi N, Takano H. Size variation of pollen grains in some *Rubus* species. J Phytogeogr Taxon. 1980; 28: 27–32.

37. Kosenko VN, Nguen TH, Jacovlev GP. Palynomorphological study of the representatives of the genus *Rubus* (Rosaceae) in the flora of Vietnam. Bot Z. 1982; 69: 497–503.

38. Fedoronchuk MM, Savitsky VD. Comparativeand morphological analysis of pollen for genera of the family Rosaceae Juss. of the Ukrainian flora. Ukr Bot Z. 1987; 44: 32–38.

39. Hebda RJ, Chinnappa CC. Studies on pollen morphology of Rosaceae in Canada. Rev Palaeobot Palynol. 1990; 64: 103–108.

40. Hebda RJ, Chinnappa CC. Studies on pollen morphology of Rosaceae. Bot Lett. 1994; 141: 183–193.

41. Ueda Y. 1992. Pollen surface morphology in the genus *Rosa*, related genera. Jpn J Palynol. 1992; 38: 94–105.

42. Tomlik-Wyremblewska A. Pollen morphology of genus *Rubus* L. Part II. Introductory studies on the Malesian species of subgenus *Micranthobatus* Fritsch. Acta Soc Bot Pol Pol Tow Bot. 2000; 69: 31–40.

43. Li WL, He SA, Gu Y, Shu P, Pu ZM. Pollen morphology of the genus *Rubus* from China. Acta Phytotax. Sin. 2001; 39: 234–247.

44. Tomlik-Wyremblewska A, Van der Ham RWJM, Kosiński P. Pollen morphology of genus *Rubus* L. Part III. Studies on the Malesian species of subgenera *Chamaebatus* L. and *Idaeobatus* L. Acta Soc Bot Pol Pol Tow Bot. 2004; 73: 207–227.

45. Wang XR, Tang HR, Huang LH, Zong ZD, Xiao LF, Hua QD, et al. Comparative studies on pollen submicroscopic morphology of some wild species and cultivars of bramble (*Rubus* L.). Yuan Yi Xue Bao. 2007; 34: 1395–1404.

46. Ghosh R, Paruya DK, Acharya K, Ghoraid N, Bera S. How reliable are non-pollen palynomorphs in tracing vegetation changes and grazing activities? Study from the Darjeeling Himalaya, India. Palaeogeogr Palaeoclimatol Palaeoecol. 2017; 475: 23–40. https://doi.org/10.1016/j.palaeo.2017.03.006

47. Gupta C, Dash SS. A new species of *Rubus* (Rosaceae) from Arunachal Pradesh, India. Blumea 2018; 63: 26–30. https://doi.org/10.3767/blumea.2018.63.01.04

48. Motyleva S, Gruner L, Semenova L. The morphology of pollen grains of some cultivars *Rubus fruticosus* L. Agrobiodiversity for Improving Nutrition, Health and Life Quality 2018; 2: 1–6. https://doi.org/10.15414/agrobiodiversity.2018.2585-8246.001-006

49. Erdtman G. The acetolysis method. A revised description. Sven Bot Tidskr. 1960; 54: 561–564.

50. Erdtman G. Pollen morphology and plant taxonomy. Angiosperms. An introduction to palynology. Stockholm: Almquist and Wiksell; 1952.

51. Punt W, Hoen PP, Blackmore S, Nilsson S, Le Thomas A. Glossary of pollen and spore terminology. Rev Palaeobot Palynol. 2007; 1431: 1–81. https://doi.org/10.1016/j.revpalbo.2006.06.008

52. Halbritter H, Hess Ulrich S, Grímsson F, Weber M, Zetter R, Hesse M., et al. Illustrated Pollen Terminology. 2nd ed. Vienna: Springer; 2018.

53. Shapiro SS, Wilk MB. An analysis of variance test for normality (complete samples). Biometrika 1965; 52: 591–611.

54. Rencher AC. Interpretation of canonical discriminant functions, canonical variates, and principal components. Am Stat. 1992; 46: 217–225.

55. Seidler-Łożykowska K, Bocianowski J. Evaluation of variability of morphological traits of selected caraway (*Carum carvi* L.) genotypes. Ind Crops Prod. 2012; 35: 140–145. https://doi.org/10.1016/j.indcrop.2011.06.026

56. Camussi A, Ottaviano E, Caliński T, Kaczmarek Z. Genetic distances based on quantitative traits. Genetics 1985; 111: 945–962.

57. Mahalanobis PC. 1936. On the generalized distance in statistics. Proc Natl Acad Sci India A. 1936; 12: 49–55.

58. Candau P, Romanos LG. Atlas polínico de Andalucía Occidental. Sevilla: Instituto de Desarrollo Regional de la Universidad de Sevilla; 1987. pp. 179–184.

59. Ghosh A, Saha I. Pollen morphological study of some selected Indian taxa of Rosaceae. Indian J Applied & Pure Bio. 2017; 32: 121–130.

60. Ueda Y, Tomita H. Morphometric analysis of pollen patterns in Roses. Hort J. 1989; 58: 211–220.

61. Ueda Y, Okada Y. Discrimination of rose cultivar groups by pollen surface structure. J Hortic Sci. 1994; 69: 601–607.

62. Polyakova TA, Gataulina GN. Morphology and variability of pollen of the genus *Spiraea* L. (Rosaceae) in Siberia and the Far East. Contemp Probl Ecol. 2008; 1: 420–424. https://doi.org/10.1134/S199542550804005X

63. Wrońska-Pilarek D, Jagodziński AM. Systematic importance of pollen morphological features of selected species from the genus *Rosa* (Rosaceae). Plant Syst Evol. 2011; 295: 55–72. https://doi.org/55-72.10.1007/s00606-011-0462-y

64. Wrońska-Pilarek D, Bocianowski J, Jagodziński AM. Comparison of pollen grain morphological features of selected species of the genus *Crataegus* L. (Rosaceae) and their spontaneous, interspecific hybrids. Bot J Linn Soc. 2013; 172: 555–571. https://doi.org/10.1111/boj.12033

